# Chromosome-level assemblies of the *Pieris mannii* butterfly genome suggest Z-origin and rapid evolution of the W chromosome

**DOI:** 10.1101/2023.03.29.534694

**Authors:** Lucas A. Blattner, Simona Ruffener, Daniel Berner

## Abstract

The insect order Lepidoptera (butterflies and moth) represents the largest group of organisms with ZW/ZZ sex determination. While the origin of the Z chromosome predates the evolution of the Lepidoptera, the W chromosome is considered younger, but its origin is debated. To shed light on the origin of the lepidopteran W, we here produce chromosome-level genome assemblies for the butterfly *Pieris mannii*, and compare the sex chromosomes within and between *P. mannii* and its sister species *P. rapae*. Our analyses clearly indicate a common origin of the W chromosomes of the two *Pieris* species, and reveal similarity between the Z and W in chromosome sequence and structure. This supports the view that the W originates from Z-autosome fusion rather than from a redundant B chromosome. We further demonstrate the extremely rapid evolution of the W relative to the other chromosomes and argue that this may preclude reliable conclusions about the origin of the W based on comparisons among distantly related Lepidoptera. Finally, we find that sequence similarity between the Z and W chromosomes is greatest toward the chromosome ends, perhaps reflecting selection for the maintenance of recognition sites essential to chromosome segregation. Our study highlights the utility of long-read sequencing technology for illuminating chromosome evolution.

**Significance:** Lepidoptera (butterflies and moths) typically exhibit a ZW/ZZ sex determination system, but the origin of the W chromosome is controversial. Based on a chromosome-level reference genome for the Southern Small white butterfly and comparative genomic analyses, we propose that the W chromosome in this group of butterflies derives from the Z chromosome and evolves extremely rapidly.

## Introduction

In many organisms, individuals are either male or female throughout their life, with sex being determined by a genetic factor located on a chromosome (Bachtrog et al. 2014; Blackmon et al. 2017; Furman et al. 2020). Depending on whether males or females produce different gametes for this sex chromosome, male-heterogametic systems (XY/XX, or X-/XX if the Y chromosome has been lost in males) are distinguished from female-heterogametic systems (ZW/ZZ or Z-/ZZ). The largest known female-heterogametic group is the Lepidoptera (Blackmon et al. 2017), the insect order including moths and butterflies. Because evolutionarily basal Lepidoptera exhibit a Z-/ZZ system and share this feature with their sister order, the Trichoptera (caddisflies), this sex chromosome configuration is considered the ancestral state in Lepidoptera (Traut & Marec 1996; Lukhtanov 2000; Traut et al. 2008; Sahara et al. 2012). However, phylogenetically more ‘advanced’ Lepidoptera, the so-called Ditrysia clade including around 99% of all extant Lepidopera species (Mutanen et al. 2010), generally show a ZW/ZZ system (including Z-/ZZ caused by secondary losses of the W chromosome). Since the Z chromosome is conserved across basal and more advanced Lepidoptera (Lukhtanov 2000; Sahara et al. 2012; Fraïsse et al. 2017), this implies that within the lepidopteran lineage, the W chromosome must have arisen long after the evolution of the Z chromosome. The specific mechanism through which the W originated, however, remains controversial.

One hypothesis postulates that the W chromosome of Lepidoptera arose from a fusion of the Z chromosome with an autosome (Lukhtanov 2000; Traut et al. 2008) (fig. 1). This scenario is typically referred to as a Z-origin of the W and we adopt this wording, noting that technically the W chromosome here derives from an autosome. Such fusions may be advantageous if autosomes harbor sexually antagonistic polymorphisms (Blackmon et al. 2017) and may appear particularly feasible in the Lepidoptera exhibiting holocentric chromosomes. That is, lepidopteran chromosomes have centromere activity along their entire length (as opposed to a single point centromere), making segregation during meiosis relatively robust to chromosome fusions and fissions (Luktanov et al. 2018; Senaratne et al. 2022). An alternative hypothesis holds that the lepidopteran W derives from a supernumerary (non-essential) B chromosome recruited for female-specific performance and that started to segregate with the Z chromosome (Lukhtanov 2000; Dalikova et al. 2017; Fraïsse et al. 2017; Lewis et al. 2021). Evaluating these two main ideas on the origin of the lepidopteran W chromosome has been challenging because the W generally contains few genes and is crammed with repeated sequences (Abe et al. 2005a; Sahara et al. 2012; Traut et al. 2013; Lewis et al. 2021). This complicates W chromosome sequence assembly and sequence-based comparative inference.

**Fig. 1.**
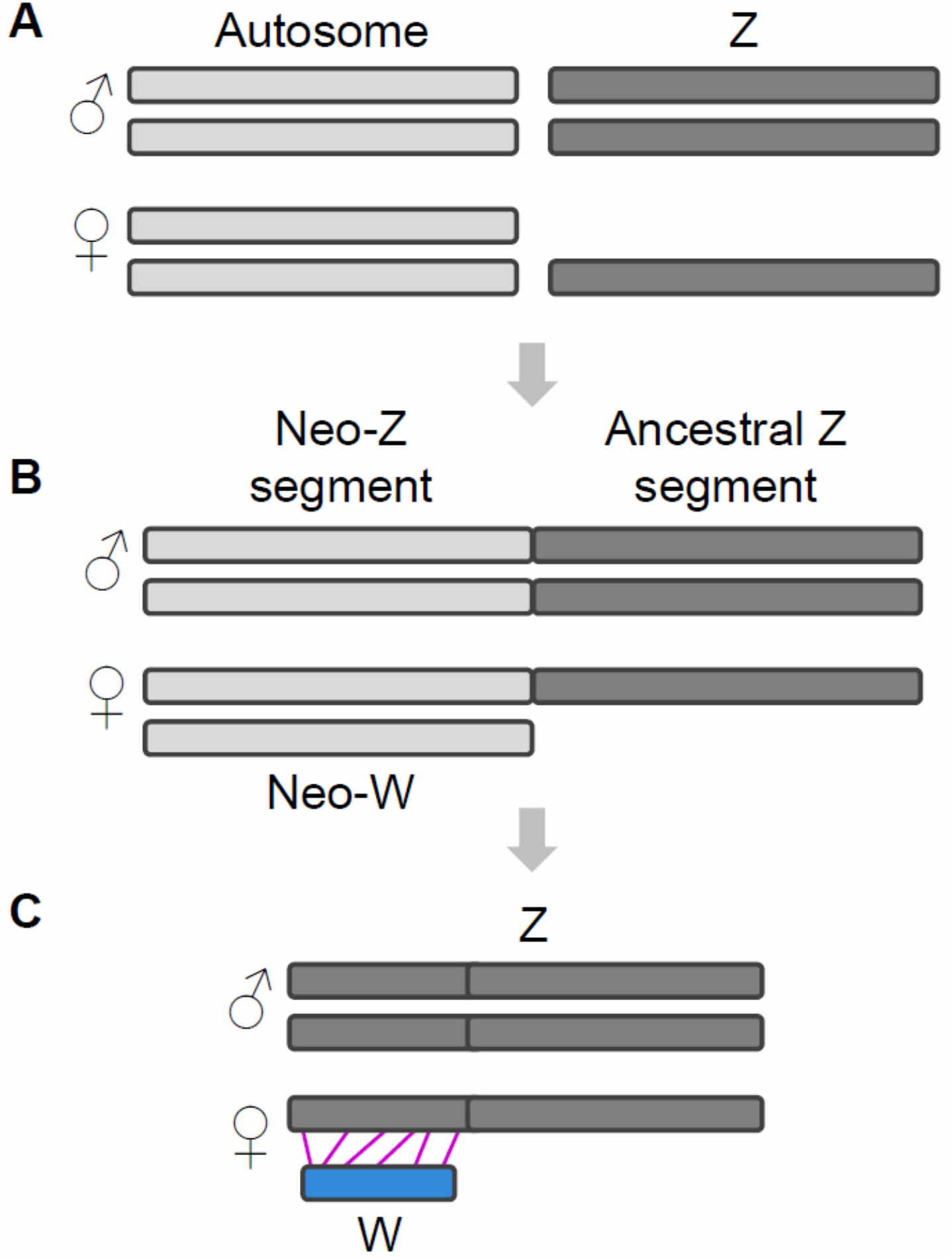
Possible evolutionary pathway from an ancestral Z-/ZZ sex chromosome configuration (A) to a ZW/ZZ system through the fusion of an autosome with the ancestral Z chromosome (B). (C) After extensive evolution of the sex chromosomes originating from the fusion, the W may retain some homology to formerly autosomal segments of the Z (indicated by purple lines).

Progress in long-read sequencing technology, however, promises to offer new insights into the origin of the lepidopteran W chromosome. In this study, we assemble the genome of a female and a male of the Southern Small White butterfly *Pieris mannii* (Mayer, 1851), including the W, from long-read sequence data. Our first goal is to use these assemblies, and those of other Lepidoptera species for which the W chromosome sequence is available, to explore the origin of the W chromosome. Our second goal is to use our assemblies to characterize the similarity between the Z and W chromosomes. The motivation for the latter is that in the Lepidoptera, recombination occurs in the homogametic males only (Traut et al. 2008). Hence the W chromosome is recombinationally isolated, a condition favoring mutational sequence degeneration (Charlesworth 1996; Bachtrog 2013), including gene loss and the accumulation of repeated sequences. Despite the expected divergence of the W from the Z, some sequence similarity between these sex chromosomes is expected to be conserved to ensure their faithful pairing and segregation during meiosis (Traut et al. 2008), but direct sequence-based evidence on these requirements is needed.

## Results

### Chromosome-level genome assemblies for Pieris mannii

A wild male and female adult of *P. mannii* were caught in 2021 near Basel, Switzerland. High molecular weight DNA was extracted from both individuals and PacBio sequenced in circular consensus (CCS) mode. Assembling the resulting HiFi reads yielded pseudohaploid raw assemblies, hereafter called the v1 assemblies, consisting of 111 (male) and 103 (female) contigs and displaying a total length of around 300 megabases (Mb) (table 1). N50 values exceeded 11 Mb, and benchmarking using single-copy orthologous genes (BUSCO; Simao et al. 2015) from Lepidoptera indicated >99% completeness. With this contiguity and completeness, our v1 assemblies compare well to the highest-quality butterfly genomes currently available (Ellis et al. 2021).

**Table 1.** Summary statistics describing the male and female Pieris mannii genome assemblies. BUSCO values are given for analyses performed with the Lepidoptera database, and the insect database in parentheses.

| Assembly | Male v1 | Male v2 | Female v1 | Female v2 |
| --- | --- | --- | --- | --- |
| Number of contigs/chromosomes | 111 | 25 | 103 | 26 |
| Total assembly length (Mb) | 296.4 | 290.9 | 302.8 | 293.3 |
| % BUSCOs | 99.3 (98.9) | 99.3 (98.9) | 99.1 (98.8) | 99.1 (98.9) |
| N50 | 12.12 | 12.21 | 11.79 | 12.19 |

Aligning the v1 assemblies to a high-quality genome (Lohse et al. 2021) of the closely related sister species *Pieris rapae* (divergence time around 3.3 million years; Wiemers et al. 2020) revealed that the vast majority of the 25 *P. mannii* chromosomes, as determined cytogenetically (Lorkovic 1985; this excludes the W chromosome), were already represented in full length by contigs of the v1 assemblies (fig. S1; table S1). Only a few autosomes were fragmented due to insertion-deletion heterozygosity in repeat-rich regions and required scaffolding by merging 2-3 contigs. Almost all chromosomes exhibited on both ends the standard (TTAGG)n repeats typical of insect telomeres (Okazaki et al. 1993) (table S1). Beyond the contigs homologous to *P. rapae* chromosomes (or chromosome segments), the remaining contigs were generally < 100 kilobases (kb) long. A notable exception was a 2.17 Mb circular contig identified in both the male and female assembly, for which BLAST search against the full GenBank nucleotide collection indicated close similarity to *Spiroplasma phoeniceum* (GenBank accession CP031088.1). *Spiroplasma* bacteria are often found in insect tissues (Hackett & Clark 1989; Ammar & Hogenhout 2006), hence this contig likely derives from such an endosymbiont of P. mannii.

To obtain a clean *P. mannii* male and female reference genome, these short contigs were excluded, retaining only the actual chromosomes plus the mitochondrial DNA sequence assembled separately for each sex. These final v2 genomes accounted for 98.1 and 96.8% of the male and female v1 assemblies. Discarding the minor contigs did not reduce BUSCO scores (table 1), hence we consider the v2 assemblies highly complete chromosome-level genomes. Chromosome lengths proved highly consistent between the male and female v2 builds (fig. 2A). Interestingly, however, all autosomes were longer (16% on average) in the *P. mannii* assemblies than in the *P. rapae genome* (fig. 2B), suggesting divergence through genome contraction/expansion between the sister species. For consistency, the *P. mannii* chromosomes in the v2 genomes are numbered according to their homologues in *P. rapae* (Lohse et al. 2021). Both v2 assemblies were subjected to *ab initio* gene prediction, and functional annotation identified 14,010 (UniProt/Swiss-Prot) and 9,769 (UniProt/TrEMBL) protein coding sequences for the male, and 14,183 and 10,369 genes for the female.

**Fig. 2.**
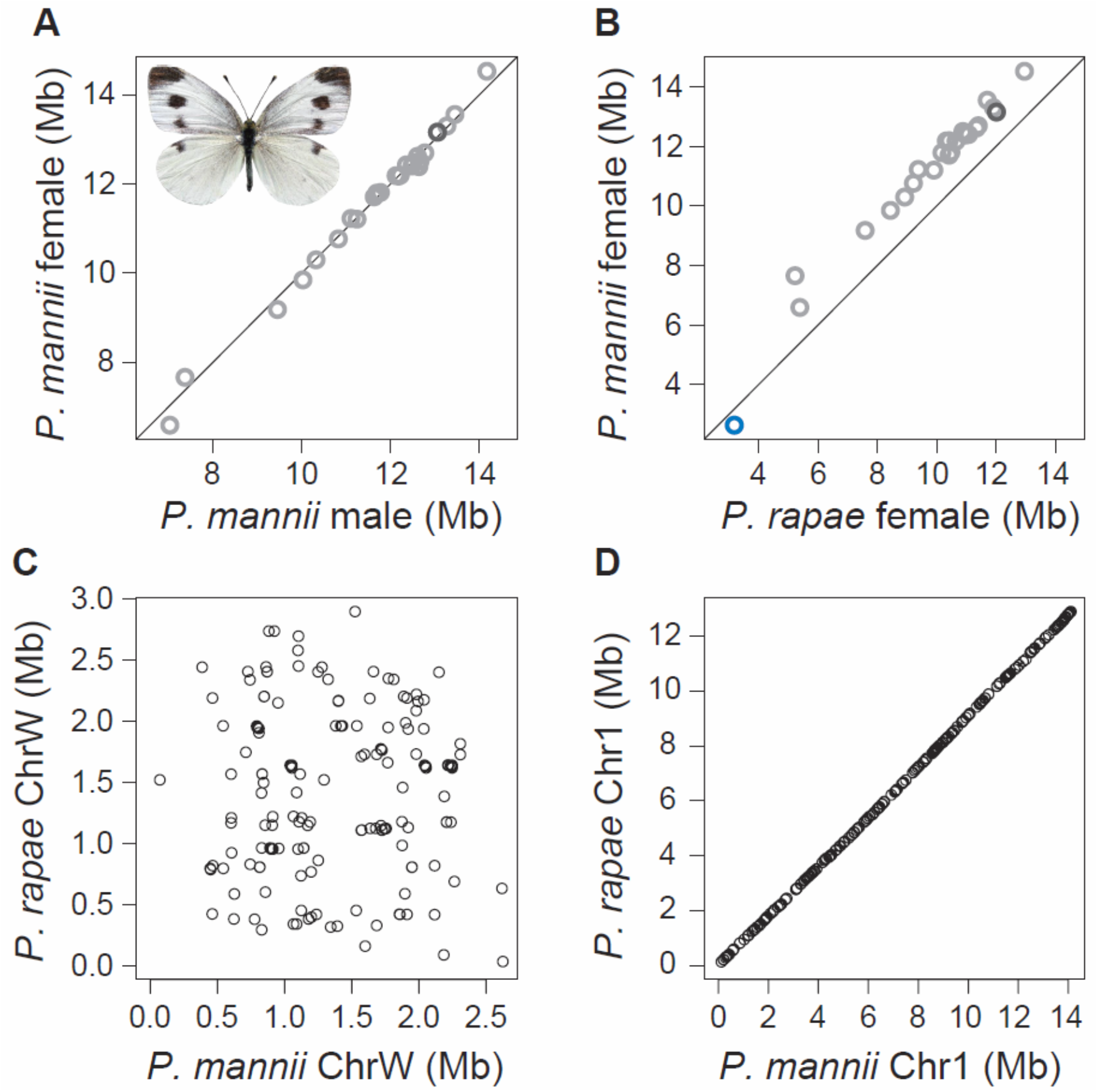
Comparison of individual chromosome lengths between (A) the *Pieris mannii* male and female v2 genome assemblies, and (B) the female *P. mannii* and *P. rapae* assemblies. Autosomes are shown in light gray, the Z chromosome in dark gray, and the W chromosome in blue. In (C), the positions of sequence tags from the *P. mannii* W chromosome (not repeat-masked) are plotted against their corresponding alignment positions on the *P. rapae* W chromosome (n = 226 tags with unique alignment). An analogous analysis based on 226 sequence tags drawn at random from a representative autosome (chromosome 1) is shown in (D).

### Identification of the P. mannii sex chromosomes

The male and female *P. mannii* Z chromosomes (13 Mb long) were identified by mapping the Z chromosome of *P. rapae* to each v1 assembly (fig. S1). We then confirmed this identification based on a comparison of read depth between the sexes (Roesti et al. 2013; Fraisse et al. 2017; Wan et al. 2019), expecting that the heterogametic ZW females exhibited only half the read depth of the homogametic ZZ males across Z chromosome segments lacking homology with the W. Aligning RAD sequence data from 19 individuals per sex to the male v2 genome, we indeed found the predicted relative reduction in female read depth across most of the Z chromosome, a pattern not observed on any autosome (fig. 3, top).

**Fig. 3.**
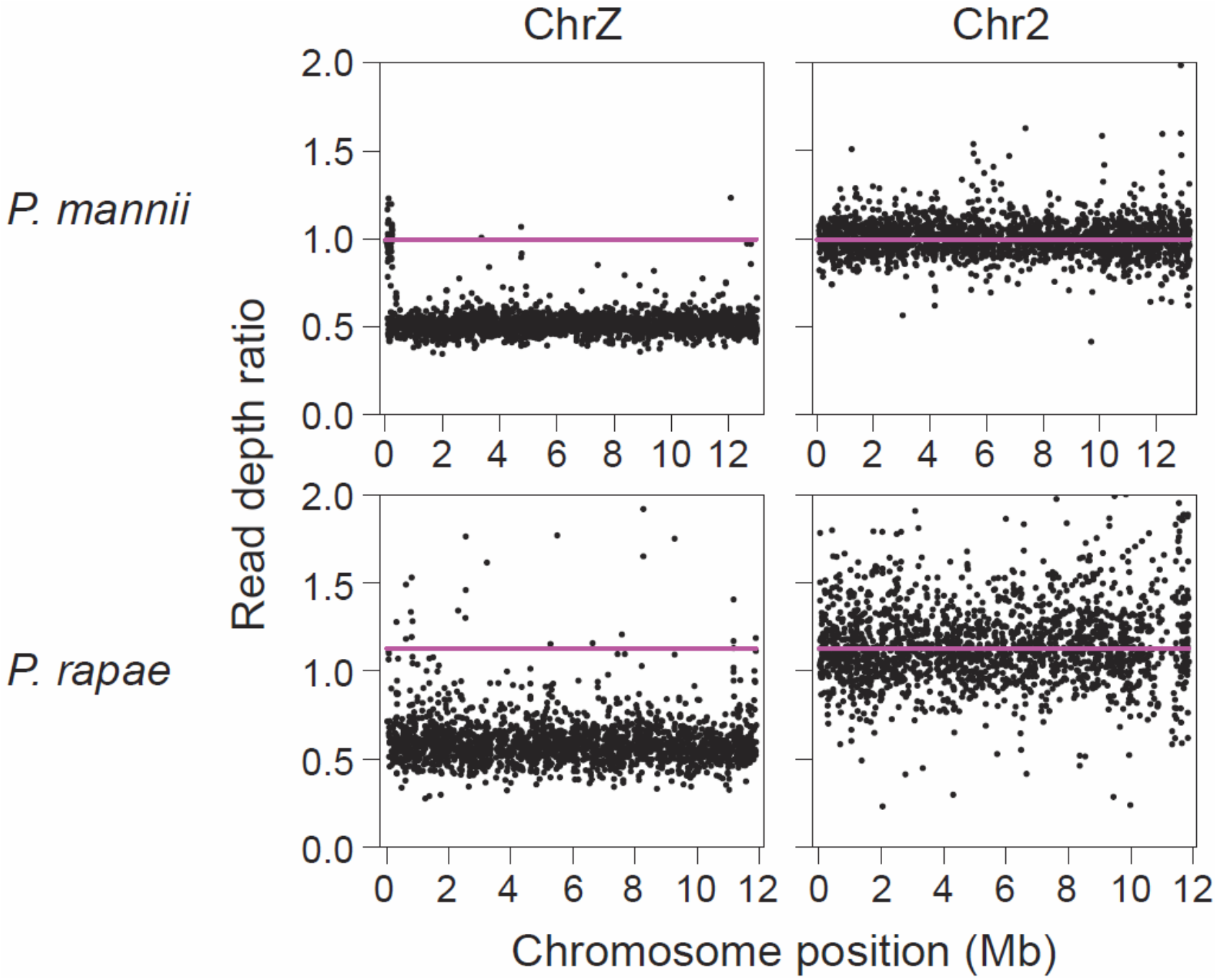
Female to male RAD locus read depth ratio along the Z chromosome, and along an exemplary autosome of approximately similar length, for *P. mannii* and *P. rapae*. RAD loci with a balanced read depth between the sexes (ratio around 1) reflect sequences present on the two parental chromosomes in each sex. By contrast, a reduced read depth in females (ratio around 0.5) along most of the Z chromosome indicates segments hemizygous in females, hence missing on the W chromosome. The purple horizontal line represents the average read depth ratio observed across the autosomes. The number of individuals is 19 and 6 per sex for *P. mannii* and *P. rapae*, and the number of RAD loci on the Z and chromosome 2 is 2501 and 2138 for *P. mannii*, and 2057 and 1817 for *P. rapae*.

A candidate *P. mannii* W chromosome, 2.6 Mb long and exhibiting telomere repeats on one end, was identified by aligning the sister species’ W chromosome (derived from a female) to our female v1 assembly (fig. S1), and confirmed by read depth analysis. For the latter, we predicted the presence of female-limited RAD loci along this contig. In line with this prediction, we observed 13 RAD loci (characterized in table S2) spread across the candidate W contig consistently present in all 19 females, but completely lacking in all 19 males. Considering that lepidopteran W chromosomes are generally enriched for repeated sequences (Traut et al. 2008; Sahara et al. 2012), we next repeat-masked the genome and found a much lower proportion (around 20% only) of non-repeated DNA on the candidate W contig compared to the autosomes and the Z (> 60%, fig. 4, left; details on the relative importance of different repeat categories across all chromosomes are given for both *P. mannii* and *P. rapae* in fig. S2). Correspondingly, we expected poor unique alignment success of W-derived sequences compared to autosomal or Z-derived sequences, hence a reduced density of RAD loci along the candidate W, which was confirmed (fig. 4, middle). Finally, the candidate W contig exhibited an unusually high GC content (fig. 4, right), which appears to be a characteristic feature of lepidopteran W chromosomes (Wan et al. 2019; Lewis et al. 2021; Lohse et al. 2021). Taken together, these analyses made clear that the female v1 assembly contained the W chromosome, which was retained for the female v2 assembly (fig. 2B; table S1).

**Fig. 4.**
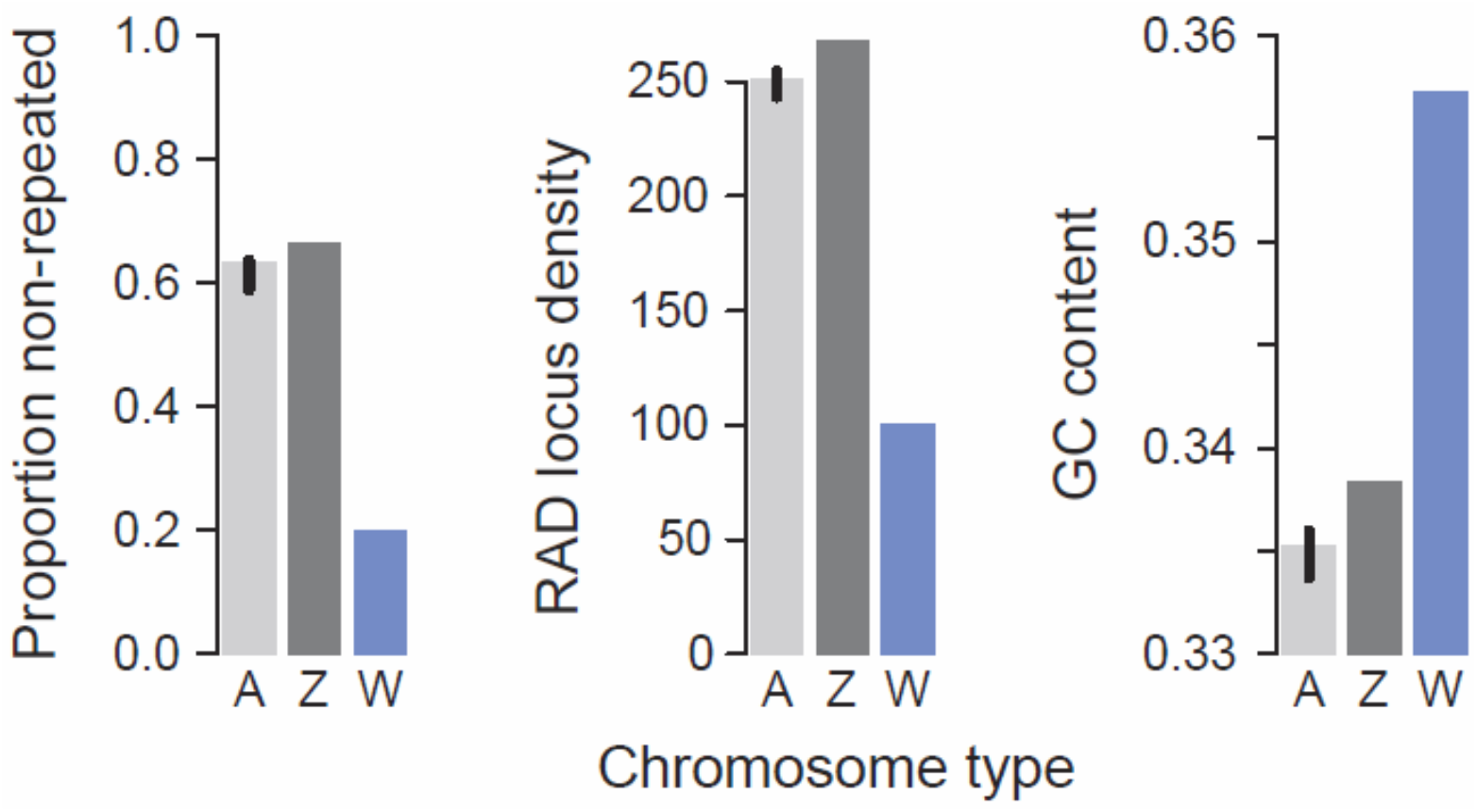
Distinctive signatures of the *P. mannii* W chromosome include an exceptionally low proportion of non-repeated DNA, a low density of loci to which RAD sequences align uniquely (expressed as loci per Mb), and a high GC content, relative to the autosomes (A) and the Z chromosome. The autosomal values are the medians across all 24 autosomes, with the error bars representing the associated 95% bootstrap compatibility intervals.

### Sequence similarity between the W and Z chromosomes

To shed light on the potential origin of the W chromosome, we extracted short (150 bp) contiguous sequences from the *P. mannii* W chromosome, and aligned these ‘W sequence tags’ to multiple target genomes. Based on the resulting alignments, we then determined ‘alignment density’, defined as the number of uniquely aligned W sequence tags per Mb, for each target chromosome. To improve the interpretability of chromosome sequence similarity as quantified by alignment density, the sequence tags (n = 3023) were extracted only from those W chromosome segments remaining after hard repeat masking, although qualitatively similar results were obtained with sequence tags from the non-masked W.

The W sequence tags were first aligned to the *P. mannii* male genome (lacking the W). This revealed that the Z chromosome displayed a relatively high alignment density, although two autosomes showed a slightly higher alignment density (fig. 5, top left). We also observed alignment to the mitochondrial DNA: around 4% of the sequence tags were identified as mitochondria-derived, including matches to Cytochrome-c-Oxidase (all three subunits) and two other genes, thus indicating the presence of nuclear mitochondrial DNA segments (NUMTs). We then aligned the *P. mannii* W sequence tags to the *P. rapae* genome and found the highest alignment density for the W chromosome (fig. 5, bottom left).

**Fig. 5.**
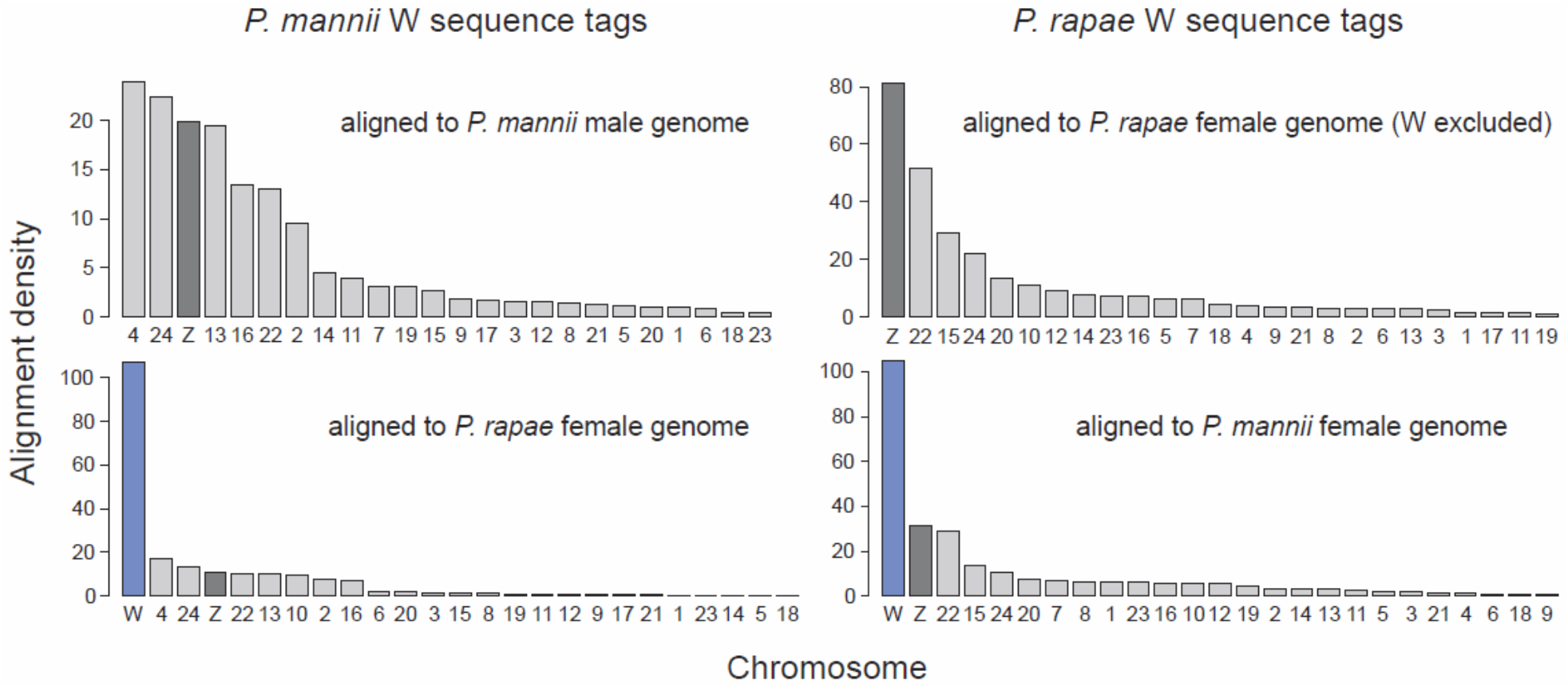
Exploring W chromosome similarity based on the alignment of sequence tags within and between *Pieris* species. The upper row shows the chromosome-specific density (alignments per Mb) of *P. mannii* and *P. rapae* sequence tags extracted from the repeat-masked W chromosome when aligned to the conspecific genome lacking the W (in *P. mannii* the male genome, in *P. rapae* the female genome with the W chromosome excluded). The lower row shows the alignment density of the same W sequence tags aligned to the female genome (including the W) of the sister species. Autosomes are plotted in light gray, Z chromosomes in dark gray, and W chromosomes in blue.

As a next step, we repeated the above analyses by using sequence tags (n = 5203) extracted from the repeat-masked W chromosome of *P. rapae*. Aligning these against the *P. rapae* genome with the W chromosome excluded, we found that the Z clearly had the highest alignment density among all chromosomes (fig. 5, top right). Aligning the *P. rapae* W sequence tags to the female genome of its sister species again produced the highest alignment density for the W (fig. 5, bottom right). Taken together, these explorations of sequence similarity based on W sequence tags within the genus *Pieris* leave little doubt that the W chromosomes of *P. mannii* and *P. rapae* originate from a shared ancestor. This view is in line with the broad-scale chromosome alignment between the sister species performed during genome assembly, indicating clear sequence homology between the W chromosomes (fig. S1, ‘ChrW’ in ‘Female’ section). Moreover, the high Z-W sequence similarity observed within both species provides a first indication that the W chromosome originates from the Z.

To explore potential sex chromosome similarity on a deeper time scale, the W sequence tags of both *Pieris* species were also aligned to three available Lepidoptera genomes containing assembled and identified Z and W chromosomes (*Dryas iulia*, Nymphalidae, Lewis et al. 2021; *Kallima inachus*, Nymphalidae, Yang et al. 2020; *Spodoptera exigua*, Noctuidae, Zhang et al. 2019). These species are all members of the Ditrysia (i.e., advanced Lepidoptera), and started diverging from *Pieris* around 100 million years ago. In these analyses between lepidopteran families, the unique alignment success of W chromosome sequence tags across all target chromosomes together ranged between 0.02% and 1.6% only. This is extremely low compared to > 40% unique alignment success in both between-species analyses within the genus *Pieris*. Furthermore, a qualitative characterization of 16 *P. mannii* sequence tags aligning uniquely to at least two of the three distantly related species using protein similarity search produced matches only for a mitochondrial gene and a transposon-related Pol polyprotein, thus suggesting that sequence similarity among the genera considered may mostly concern transposable elements. Given these indications of vast sequence divergence between the *Pieris* W chromosomes and the entire genomes of the distantly related Lepidoptera, we refrained from drawing conclusions about the origin of the W chromosome beyond the more recently split *Pieris* species.

### Structural similarity between the W and Z chromosomes

Having found sequence similarity among sex chromosomes within and between *Pieris* species, we next aimed to characterize to what extent the W and Z chromosomes within each species also retained similarity in physical structure. In *P. mannii*, a first noteworthy pattern emerged from the RAD sequence-based read depth analysis. We here observed a narrow (254 kb) region in the left periphery of the Z chromosome exhibiting numerous RAD loci with a balanced read depth between the sexes (fig. 3, top left), the standard read depth ratio seen throughout the autosomes. As the RAD data were aligned to the male assembly, this pattern must arise from the RAD loci in this Z chromosome region having a homologous counterpart on the W. However, aligning the RAD sequences from these loci to the hard-masked W chromosome did not produce a single alignment. This suggests that this region of the Z chromosome harbors sequences repeated on the W, thus making conclusions about structural similarity between the sex chromosomes based on read depth difficult.

More robust information on the structural similarity between the *P. mannii* sex chromosomes, however, emerged from calculating the correlation between the position of the W sequence tags on the repeat-masked W and their alignment position on the other chromosomes. This positional correlation was nearly perfect for the Z chromosome (Spearman’s rank correlation *r*_S_ = 0.998), but lower for the autosomes (*r*_S_ always <= 0.66) (fig. 6, top left), indicating particularly high similarity in chromosome structure between the Z and W. Plotting the W against the Z positions for all W sequence tags mapping to the Z explained this similarity: the W chromosome broadly mirrored a segment located on the right periphery of the Z chromosome (fig. 6, top right). Overall, 97% of the W sequence tags aligning to the Z chromosome mapped to the peripheral 1 Mb on the right side of the Z. A closer look into this region further revealed multiple windows across which the Z and W were collinear, a structural association obtained irrespective of whether the W chromosome was repeat-masked when extracting the sequence tags (fig. 7A). The latter was in striking contrast to the three autosomes also exhibiting a relatively high alignment density (i.e., the chromosomes 4, 24, 13; fig. 5, top left). For chromosome 4, for instance, sequence tags generated without repeat masking the W showed that similarity in sequence and structure was driven strongly by a single short (81.6 kb) region copied from this autosome into the W chromosome, where this segment expanded to a 200 kb region through extensive tandem repeat multiplication (fig. 7B). Among all W sequence tags aligning to chromosome 4, 92% mapped to this single region. Similar evidence of translocations followed by repeat expansion also emerged for the chromosomes 24 and 13 (fig. S3).

**Fig. 6.**
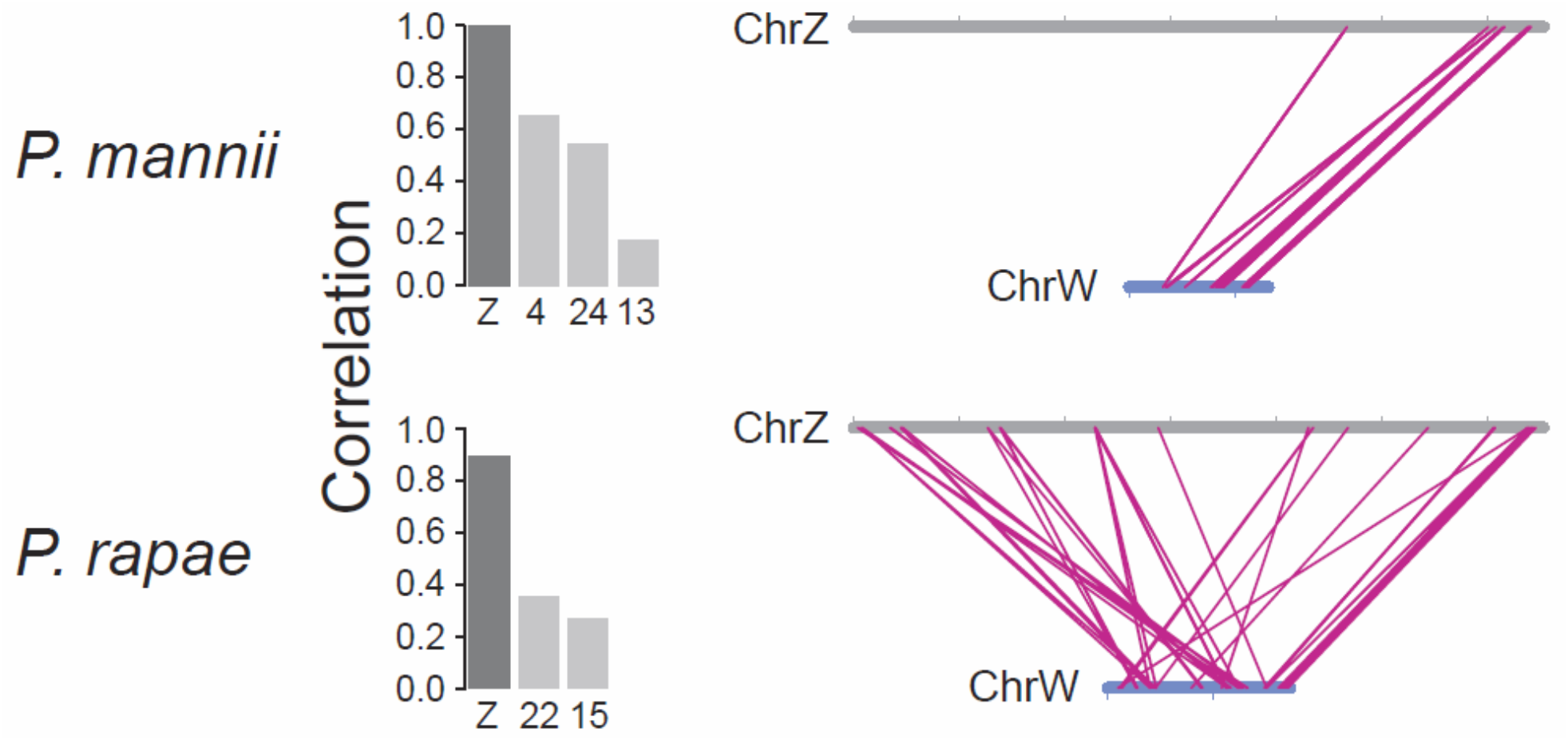
Structural similarity between the Z and W chromosomes in *P. mannii* and *P. rapae*. The left graphs show the Spearman coefficients (*r*_S_) for the correlations between W sequence tag positions on their source chromosome (i.e., the repeat-masked W), and on all target chromosomes to which at least 120 tags aligned uniquely. The Z chromosome is shaded dark gray, autosomes light gray. In the right graphs, the positions of W sequence tags are connected to their alignment positions on the Z chromosome. The sequence tags were extracted from the repeat-masked W, and were required to map uniquely to the Z when aligning them to the genome lacking the W (in *P. mannii* the male genome, in *P. rapae* the female genome with the W chromosome excluded). Sample size is n = 260 and 979 sequence tags for *P. mannii* and *P. rapae*. The sex chromosomes are drawn to scale, the tick marks delimit 2 Mb intervals. The *P. rapae* W chromosome was reversed relative to its orientation in the reference genome.

**Fig. 7.**
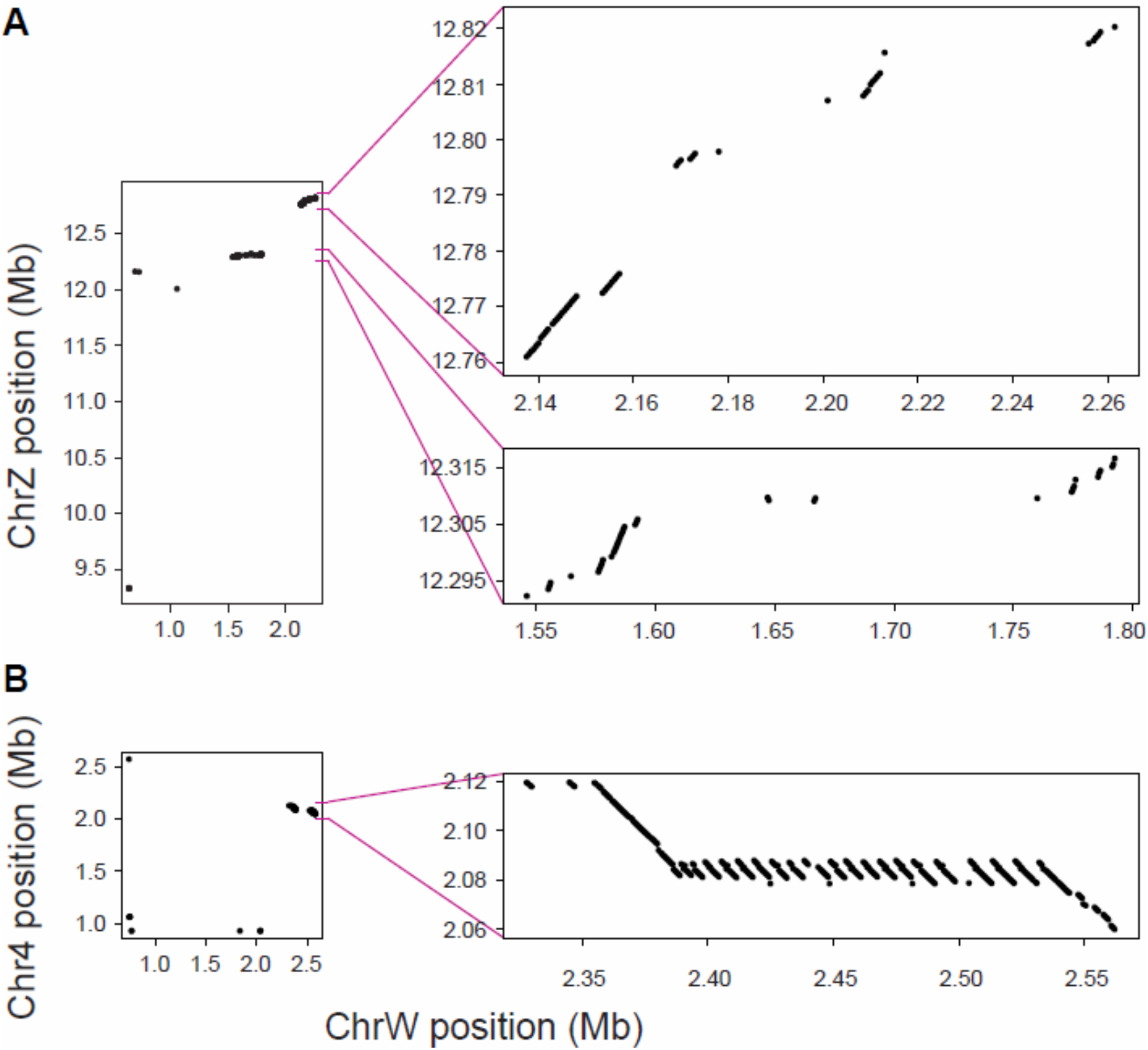
(A) Physical position of all *P. mannii* W chromosome sequence tags aligning to the Z chromosome. The left graph is based on sequence tags derived from the repeat-masked W (hence uses the same data as in fig. 6, top right). The right graphs represent close-ups of the two regions showing extended Z-W sequence similarity, based on sequence tags extracted from the W chromosome *not* repeat-masked. In (B), analogous data are shown for chromosome 4.

Taken together, these analyses based on W sequence tags make clear that the *P. mannii* W chromosome retains exceptional structural similarity to the Z chromosome, and that this similarity is largely restricted to one periphery of the Z. Moreover, some autosomes displaying a relatively high sequence similarity to the W appear to do so because of translocations into the W, followed by repeat expansion. The only substantial similarity in chromosome structure not obviously associated with repeated DNA is between the Z and W. We thus propose that the similarity in both sequence content and structure between the sex chromosomes of *P. mannii* reflects remnant homology persisting in the face of rapid W chromosome evolution due to insertion, deletion, and repeat mutations.

Repeating these explorations of Z-W structural similarity in *P. rapae*, we found again that among all the chromosomes, the Z displayed the strongest correlation with the W in the position of W sequence tags (*r*_S_ = 0.89; all autosomes *r*_S_ <= 0.35; fig. 6, bottom left). The Z-W homology captured by the W sequence tags in *P. rapae* was not as strongly biased toward one single chromosome periphery as in *P. mannii* (fig. 6, bottom right), but the peripheral 1 Mb on the right side of the Z chromosome still accounted for 62% of the W sequence tags aligning to the Z (87% of the W sequence tags came from the peripheral 1 Mb on either side of the Z). The difference in the structural Z-W similarity between the sister species suggests substantial divergence between the W chromosomes. Supporting this idea, the positions of *P. mannii* W chromosome sequence tags proved largely uncorrelated (*r*_S_ = -0.033) to their mapping positions on the W of *P. rapae* (fig. 2C, based on sequence tags from the W chromosome not repeat-masked; a similar result was obtained when using sequence tags from the masked W, fig. S4). By contrast, this correlation was nearly perfect on all autosomes and the Z (fig. 2D, fig. S5B). Moreover, while the latter chromosomes were consistently longer in *P. mannii* than in *P. rapae*, the *P. mannii* W chromosome was 17% shorter than its counterpart in the sister species (fig. 2B). Recognizing the possibility of W chromosome assembly errors, the conclusion that the W chromosomes of the two *Pieris* species are dramatically more divergent than the autosomes is unlikely to be qualitatively wrong.

## Discussion

### The Pieris W chromosome likely arose from Z-autosome fusion, not from B chromosome recruitment

A key finding from our analyses based on high-quality genome assemblies relevant to our understanding of the origin of the lepidopteran W chromosome is that within both *Pieris* species, the W exhibits substantial sequence similarity to the Z, and also retains greater structural similarity to the Z than to any autosome. This is consistent with the notion that the W chromosome originates from a Z chromosome-autosome fusion (or multiple such fusions) during the early evolution of the advanced Lepidoptera (Lukhtanov 2000; Traut et al. 2008). This fusion would have turned one autosome copy into the new W chromosome and the other copy into a new segment of the Z chromosome retaining some degree of homology to the W despite the mutational degeneration of the latter (fig. 1). This view aligns with evidence of a key role of major rearrangements in the evolution of sex chromosomes in other advanced Lepidoptera (Smith et al. 2016; Mongue et al. 2017; Carabajal-Paladino et al. 2019; Hill et al. 2019; Yoshido et al. 2005, 2020; Hejnickova et al. 2021), and with the recent demonstration of a shared Z-origin of the W chromosomes in two species of Crambidae (also advanced Lepidoptera) (Dai et al. 2022).

Three alternative explanations for the observed similarity between the Z and W chromosomes, however, deserve discussion. First, the Z-W similarity may be an artifact arising from misassembly of the Z chromosome. In *P. mannii* for instance, the segment on the right periphery of the Z showing high structural similarity to the W could represent an autosomal fragment erroneously assembled into the Z and retaining homology to segments transposed from the same autosome into the W, the latter potentially originating from a B chromosome. This possibility is easily refuted; we produced two entirely independent Z chromosome assemblies for *P. mannii*, and both whole-chromosome and Z chromosome sequence tag alignments confirm that these assemblies are essentially identical (fig. S5A). Moreover, apart from minor inconsistencies in chromosome segment orientation, our Z assemblies are also nearly identical to the Z chromosome of *P. rapae* (fig. S5B).

Second, in *Pieris napi*, a congener of our main study species, a segment of the Z chromosome seems to originate from a relatively recent fusion of the broadly conserved Lepidopteran Z with an autosome homologous to chromosome 2 of the *Bombyx mori* reference genome (Hill et al. 2019). The Z-W similarity we observe could thus reflect segments of this relatively new Z region exhibiting homology to W chromosome segments also originating from that same autosome, which would again not necessarily be in conflict with a B chromosome origin of the W. This idea is also rejected: Z chromosome alignment between *P. mannii* and *P. napi* shows that the Z-W similarity uncovered in our study occurs on the Z chromosome end opposite to the more recently acquired Z region identified in *P. napi* (fig. S6).

Third, it is conceivable that the high Z-W similarity in *Pieris* simply arises from the disproportional transposition of Z chromosome segments relative to autosomal segments into a former B chromosome. Based on our data, this scenario does not appear parsimonious. The main reason is that in *P. mannii*, the alignment of sequence tags derived from the W chromosome *not* repeat-masked uncovers extensive repeated DNA around the major regions of autosome-W similarity, but not in the regions of Z-W similarity. We take this as indication that Z-W homology has not arisen from transposition, but represents true remnant homology. The main alternative to a Z-origin of the W chromosome, involving the ancient recruitment of a B chromosome (Lukhtanov 2000; Dalikova et al. 2017; Fraïsse et al. 2017; Hejnickova et al. 2019; Lewis et al. 2021), perhaps followed by its acquisition of Z chromosome sequences to facilitate segregation (see below), cannot be ruled out definitively but is not easily reconciled with our results.

### Lepidopteran W chromosomes evolve rapidly

Our data further support the notion that W chromosome evolution is extraordinarily rapid (Vitkova et al. 2007; Sahara et al. 2012; Hejnickova et al. 2021). The evolution of lepidopteran W chromosomes frequently involves sequence deletions, the incorporation of autosomal segments, and extensive sequence multiplication (Abe et al. 2005a, 2005b; Traut et al. 2013; Lewis et al. 2021; Dai et al. 2022). Our identification of autosomal segments copied into and expanding on the W chromosome in *P. mannii* exemplifies some of these processes. Clearly, the recombinationally isolated W chromosomes of the two *Pieris* sister species have diverged at a dramatically higher pace than the autosomes and the Z chromosome.

Our demonstration of vast W chromosome divergence even between relatively recently (3.3 million years) separated species has implications for the strategies used to infer the origin of the W chromosome in Lepidoptera. So far, the view that the W chromosomes have, perhaps repeatedly, arisen from B chromosomes rests on the absence of detectable Z-W homology within species, or of W-W homology among relatively deeply separated lineages (e.g., Lewis et al. 2021). Given the rapid evolution of the W, however, at least the latter may be inconclusive. In our study, the *Pieris* W sequence tags aligning successfully to the genomes of representatives of phylogenetically distant lepidopteran families were indeed so sparse that we considered inference regarding chromosome similarity unreliable. Consistent with this reservation, the only other demonstration of W-W homology between lepidopteran species so far also concerns relatively closely related species from the same family (Dai et al. 2022).

### Sequence homology in the chromosome peripheries may be required for proper Z-W chromosome segregation

Although the W has been lost in some advanced Lepidoptera (Traut et al. 2008; Yoshido et al. 2013; Hejnickova et al. 2021), indicating that the W may sometimes become dispensable, this chromosome in general likely encodes information important for female performance (Kiuchi et al. 2014; Lewis et al. 2021). The rapid evolution of the W chromosome thus raises a conundrum (Traut et al. 2008): as sequence divergence between the W and the Z progresses, homology between these chromosomes should become so low that proper pairing and segregation during meiosis are compromised. To shed light on the requirements for faithful sex chromosome segregation, we examined the physical distribution of Z-W sequence homology and found that chromosome regions retaining extensive homology were predominantly located in the peripheries of the Z chromosome. (Given its short length [fig. 2B], this pattern was not apparent on the W chromosome.) Combined with the observation of a lower W degeneration rate toward the chromosome peripheries in Pyralid moths (Vitkova et al. 2007), and with previous hints to a crucial role of the chromosome peripheries to chromosome segregation in Lepidoptera females (Rego & Marec 2003) and other organisms (reviewed in Haenel et al. 2018), we speculate that degenerating W chromosomes are selected to maintain recognition sequences homologous to the peripheries of the Z chromosomes with which they segregate.

### Conclusions

Much remains to be learned about the evolution of Lepidoptera sex chromosomes, but fresh insights will likely emerge rapidly from high-quality genome assemblies based on long-read sequencing. In this study, we generated such data for a *Pieris* butterfly. We obtained telomere-to-telomere chromosome-level assemblies by using a single wild-caught individual from each sex and relying on a single sequencing effort, highlighting the power of long-read sequencing technology. Comparing sequence content and chromosome structure based on our assemblies and that of a close congener allows us to support the idea that lepidopteran W chromosomes originate from the Z chromosome through Z-autosome fusion. Our results also highlight that lepidopteran W chromosomes evolve extremely rapidly, thus hampering the detection of shared ancestry between more deeply separated taxa. Challenges for the future include determining how frequently Z-autosome fusion has occurred across the Lepidoptera, how much Z-W sequence homology is required to maintain faithful chromosome segregation, and what segments of the W actually matter to female function. The advent of further chromosome-level genomes from across the Lepidoptera phylogeny will greatly facilitate addressing these ideas.

## Methods

### DNA extraction, long-read sequencing, and genome assembly

Multiple wild (i.e., not inbred) individuals of *Pieris mannii* were caught in July 2021 from a residential area near Basel (coordinates: 47.52069, 7.68656). An initial round of DNA extraction was performed by considering different combinations of butterfly tissues and extraction techniques, including column- and bead-based kits, and evaluated for DNA yield, purity, and integrity. The highest DNA purity and adequate yield (c. 2-5 [median: 3.6] μg DNA at a concentration of 26-60 [median: 43] ng/μL) were obtained from whole adult thorax samples, and the highest DNA integrity was achieved with phenol-chloroform-isoamyl alcohol extraction (average fragment length: 28-39 [median 38] kb). DNA from a single male and female extracted in this way was then HiFi circular consensus sequenced (CCS) as low input libraries to 40x and 30x average read depth on a PacBio Sequel IIe instrument by the Functional Genomics Center Zurich (FGCZ). Further details on DNA extraction and sequencing are provided in the Methods S1.

Pseudohaploid (i.e., not fully phased) assemblies of the HiFi reads were performed in parallel by using Hifiasm (v0.16.1; Cheng et al. 2021), HiCanu (v2.2; Nurk et al. 2020), and the IPA HiFi genome assembler (v1.5.0; Dunn & Sovic 2020). Quality metrics were calculated by using QUAST (v5.2.0; Gurevich et al. 2013), and completeness was evaluated with BUSCO (v5.4.2; Simao et al. 2015). To increase the comparability with other genome assemblies, we ran BUSCO analyses with both the Lepidoptera (lepidoptera_odb10: 16 genomes, 5286 genes) and the insect (insecta_odb10: 75 genomes, 1367 genes) lineage databases. Hifiasm clearly produced the most contiguous assemblies (table S3), so we did not consider the other assemblies for further work.

The two raw contig-level (‘v1’) assemblies directly returned from the Hifiasm assembler were subjected to homology search using the chromosome-level genome of *Pieris rapae* (Lohse et al. 2021; GenBank accession GCF_905147795.1), the sister species of *P. mannii* (Wiemers et al. 2020). For this, we mapped each of the 26 *P. rapae* chromosomes (including the W) separately against our male and female v1 assemblies using Minimap2 (v2.20; Li et al. 2018). This showed that 28 of the total 111 contigs in the male v1 assembly, and 33 of the 103 contigs in the female build, exhibited unambiguous long-range homology to *P. rapae* chromosomes (or chromosome segments) (fig. 1D; fig. S1). Closer inspection revealed that most of the *P. mannii* v1 contigs showing homology to the sister species represented chromosomes assembled full-length from telomere to telomere (table S1; fig. S1). The three non-contiguous chromosomes in the male v1 build were scaffolded with RagTag (v2.1.0; Alonge et al. 2021) by considering just the HiFi reads longer than 5 kb, and using the *P. rapae* genome as backbone. This showed that the three chromosome gaps were caused by short (24-42 bp) insertion-deletion (indel) polymorphisms located in repeat-rich regions. We fixed these gaps by incorporating the insertion haplotype. The female v1 assembly, based on slightly shallower read depth, showed seven such chromosome gaps in total, of which three could be fixed as in the male assembly by resolving indels 2-25 bp long. Four gaps, however, could not be resolved unambiguously; they were closed by incorporating sequences of 100 Ns between the properly ordered contigs. To check the robustness of the v1 W chromosome assembly, we excluded all HiFi reads shorter than 5 kb, and performed a new assembly. The W contig emerging from this assembly contained a c. 200 kb segment missing in the initial build, but was nearly identical to the original W contig otherwise; hence we treated this alternative contig as the final W chromosome. To obtain a final ‘v2’ male and female reference genome, we discarded all minor contigs from the v1 builds (95% and 73% of which were shorter than 100 kb in the male and female), and retained only the (joined) chromosomes, plus the mitochondrial genomes assembled and annotated independently from the HiFi reads by using MitoHifi (v2.2; Allio et al. 2020). All chromosomes in the v2 assemblies are ordered and named according to their homologues in the *P. rapae* genome.

To obtain repeat-masked *P. mannii* and *P. rapae* genome versions, we followed Kim & Kim (2022) and first used RepeatMasker (v4.0.9; http://repeatmasker.org) with its internal Arthropoda repeat library (Dfam v3.0) to identify known repeats. *De novo* repeat libraries specific to the assembly of each *Pieris* species were then established using RepeatModeler (v2.0.1; http://repeatmasker.org). Final hard- and soft-masked chromosomes were then obtained by another RepeatMasker run.

### Structural and functional genome annotation

The soft-masked v2 genome assemblies were structurally annotated by *ab initio* gene prediction using AUGUSTUS (v3.2.3; Stanke et al. 2006; Hoff & Stanke 2019), trained on the *P. rapae* annotation (Lohse et al. 2021). After initial training, spurious genes were removed, and training was optimized by following the recommendations for predicting genes in single genomes (Hoff & Stanke 2019). The predicted protein sequences were identified and functionally annotated with a protein-protein homology approach. Protein sequences were matched against the UniProt/Swiss-Prot database (downloaded: 2022-09-26) with the blastp algorithm implemented in BLAST+ (v2.13; Camacho et al. 2008). Considering only reads with E-values < 0.001, the protein match with the lowest E-value and the highest bitscore was transferred to the final sequence headers. Predicted protein sequences not identifiable using the high-quality Swiss-Prot database were matched against the UniProt/TrEMBL protein database and processed as described above.

### RAD sequence data and analysis

Population-level sequence data are valuable for characterizing sex chromosome similarity based on sex-specific read depth. We generated such data for *P. mannii* and *P. rapae* using restriction site-associated DNA (RAD) sequencing. We here used 38 *P. mannii* (19 per sex) and 12 *P. rapae* (6 per sex) individuals captured during the summers of 2020 and 2021 in the same region as the individuals used for the genome assemblies (table S4). DNA was extracted from adult tissue (whole thorax plus head) using the Qiagen DNeasy Blood & Tissue kit, generally obtaining concentrations of 10-50 (median 20) ng/μL. A total of 100 ng genomic DNA from each individual was digested with the PstI restriction enzyme (c. 40 k recognition sites in the *P. mannii* genome), subjected to RAD library preparation as described in Blattner et al. (2022), and paired-end sequenced to 100 or 150 bp on an Illumina HiSeq4000 or NovaSeq 6000 instrument at the Quantitative Genomics Facility, D-BSSE, ETH Zürich. The demultiplexed short reads were then filtered for those harboring the PstI restriction residual to enforce sequence homology (thus effectively reducing the data set to single-end reads), and trimmed to 91 bp if needed. To ensure relatively balanced read depth across individuals, short read files containing more that 8 million reads were reduced to that number.

The short reads were then aligned to the *P. mannii* male v2 genome (or, for the *P. rapae* individuals, the Lohse et al. 2021 genome with the Z chromosome excluded) using Novoalign (v3.00; http://www.novocraft.com/products/novoalign), allowing a total mismatch value of t400. We here chose male genomes because the presence of the W chromosome in the female genome would have caused non-unique mapping, and hence the exclusion, of sequences with Z-W similarity. The 38 resulting alignments for *P. mannii* were then uploaded together into R (R Core Team 2020) using the Rsamtools package (v2.2.1; Morgan et al. 2017), and female to male read depth ratio was calculated for all genome-wide RAD loci, provided that they were represented in at least 18 out of the 19 individuals per sex, thus ensuring high precision in the estimation of sex-specific read depth. Also, total read depth across all individuals combined had to be within 450 to 2400x, thus filtering poorly covered genome regions and obviously repeated sequences. For *P. rapae*, RAD locus representation in at least five individuals per sex, and a total read depth between 150 and 700, was required. Female-male read depth ratio at RAD loci was then plotted across the autosomes and the Z chromosome to validate the correct identification of the Z (identified by homology to the *P. rapae* Z), and to explore Z-W sequence similarity.

To validate the candidate W chromosome in *P. mannii*, as suggested by *P. rapae* sequence similarity, we aligned the short reads from all 38 individuals to the female v2 genome. We then searched for female-limited RAD loci by requiring representation in all 19 females and complete absence in all males, plus a pooled female read depth between 225 and 1200x.

Finally, we used the RAD sequence data from *P. mannii* to compare the density of high-quality RAD loci among all chromosomes. For this, we aligned the short reads from just the 19 females to the female genome. The males were excluded because they did not provide sequence information for the W chromosome. We considered all RAD loci represented in at least ten females as high-quality loci, and calculated RAD locus density as the number of such loci per Mb of sequence for each of the 26 chromosomes. For the autosomes, we estimated the 95% compatibility interval for the median density based on 10,000 bootstrap resamples.

### Analyses based on W chromosome sequence tags

To explore the similarity in sequence content and structure between chromosomes with high physical resolution, we extracted as many contiguous (non-overlapping) 150 bp segments as possible from all W chromosome segments surviving hard repeat masking, for both *Pieris* species. These sequence tags (n= 3023 and 5203 for *P. mannii* and *P. rapae*) were then converted into a fastq file for each species by adding the source position on the corresponding W chromosome for each tag, and a quality string consisting of top read quality scores (‘H’). As a robustness check, all analyses based on W sequence tags in both *Pieris* species were repeated by deriving the tags from the W chromosome without repeat masking (we here extracted 150 bp tags every 500 bp along the W; n = 5279 and 6352 for *P. mannii* and *P. rapae*). Although we consider it more difficult to infer sequence homology from the latter analyses and present only a subset, the insights from W sequence tag analyses with and without repeat masking were qualitatively similar throughout.

For both *Pieris* species, we then aligned the W sequence tags to multiple genomes by using Novoalign with a total mismatch threshold of t600. We initially aligned against each species’ own genome, but without the W chromosome (i.e., the male v2 genome for *P. mannii*, and the *P rapae* genome with the W omitted) to permit unique matches of the tags to all the other chromosomes. For each target chromosome to which at least 120 sequence tags aligned uniquely, we calculated the strength of the association in physical positions of the tags (W positions vs. positions on focal chromosome) by the Spearman correlation coefficient (r_S_; the Pearson coefficient produced qualitatively similar results). Next, we expanded the alignment analysis to between-species combinations by using a representation of available Lepidoptera reference genomes with well-assembled W chromosomes, including *Dryas iulia* (Lewis et al. 2021; GenBank accession GCA_019049465.1), *Kallima inachus* (Yang et al. 2020; Dryad dataset https://doi.org/10.5061/dryad.8w9ghx3gt), and *Spodoptera exigua* (Zhang et al. 2019; GenBank accession GCA_011316535.1). Splitting time estimates between *Pieris* and these species were obtained from TimeTree5 (Kumar et al. 2022), and the splitting time between the two *Pieris* species was obtained from Wiemers et al. (2020). Analogously to RAD locus density, the number of sequence tags aligning to a given chromosome of a target species was divided by that chromosome’s length to obtain chromosome-specific alignment density (alignments per Mb).

As robustness checks, all analyses using W sequence tags were repeated by extracting longer (300 bp) contiguous segments from the repeat-masked W chromosomes (the total alignment mismatch threshold was here adjusted to t1200). Moreover, a subset of alignments using the standard 150 bp segments were performed by halving and doubling the total mismatch threshold (i.e., t300 and t1200). Although both greater segment length and a higher mismatch threshold tended to increase alignment success, these changes always produced results consistent with the standard analyses and supporting the same conclusions (fig. S7).

### Additional explorations of chromosome similarity

To explore the collinearity between the two Z chromosome builds within *P. mannii*, we extracted sequence tags from the (not repeat-masked) male Z chromosome as described above, and aligned the tags to its female counterpart. The same approach was taken to compare a haphazardly selected autosome (chromosome 1) between *P. mannii* and *P. rapae*. Additional chromosome comparisons among *Pieris* species (including *P. napi*) involved chromosome alignment using Minimap2 and visualization in D-GENIES (Cabanettes & Klopp 2018). Unless indicated otherwise, all analyses were implemented in R (R Core Team 2020).

## Supporting information

Complete Supplementary Information

## Supplementary Material

Supplementary methods, figures, and tables (combined to a single pdf document), and the full compilation of the code underlying the study, are available at Genome Biology and Evolution online.

## Acknowledgements

We thank Nicolas Boileau for support in the wet lab; Dustin Kulanek for aiding RAD library preparation; Simon Grüter from the FGCZ for long-read sequencing advice. Computation was partly aided by Moritz Gubler and was performed at sciCORE (http://scicore.unibas.ch/) scientific computing center at the University of Basel on their HPC infrastructure. We further thank Joshua Ebner for his helpful comments on Genome Annotation, and three anonymous reviewers for constructive feedback on the initial manuscript.

## Author Contributions

Study design: DB, LB, SR; funding: DB; specimen sampling and breeding: DB, SR; wet lab work: SR, LB; data analysis: DB, LB, SR; wrote the paper: DB, SR, LB.

## Data Availability

The raw HiFi CCS reads and the v2 genome for the male and female are deposited on NCBI under the accession PRJNA885610. The Swiss-Prot and TrEMBL annotation files for both genomes and the mitochondria are available from the Dryad digital repository (doi: https://doi.org/10.5061/dryad.1vhhmgqwx). The *Spiroplasma* chromosomes from both individuals are available under NCBI BioProject accession PRJNA885610. The 50 raw demultiplexed RADseq fastq files are available from the NCBI Short Read Archive under the accession numbers listed in table S4 and NCBI BioProject accession PRJNA885610.

