## Supplementary material for "Chromosome-level assemblies of the *Pieris mannii* butterfly genome suggest Z-origin and rapid evolution of the W chromosome": Complete Supplementary Information

by Blattner et al. 2023

### Contents


- **Methods S1.** DNA extraction and sequencing p. 3
- **Fig. S1.** Correspondence between *P. rapae* chromosomes and contigs in the *P. mannii* v1 assemblies p. 6
- **Fig. S2.** Proportion of different repeat classes by chromosome for *P. mannii* and *P. rapae* p. 8
- **Fig. S3.** *P. mannii* W sequence tag positions on chrW versus chr13 and chr24 p. 9
- **Fig. S4.** *P. mannii* repeat-masked W sequence tag positions on the *P. mannii* versus *P. rapae* W chromosome p. 10
- **Fig. S5.** Check of correct assembly of the *P. mannii* Z chromosomes p. 11
- **Fig. S6.** Check of collinearity between the *P. mannii* and *P. napi* Z chromosomes p. 12
- **Fig. S7.** Robustness check for the W sequence tag analysis p. 13
- **Fig. S8.** Fragment length distribution for the male and female DNA samples submitted for long-read sequencing p. 14
- **Fig. S9.** Read length distribution and read depth for the sequenced male and female p. 15
- **Table S1.** Characterization of the chromosomes in the male and female v2 assemblies p. 16

- **Table S2.** Characterization of the 13 female-limited RAD loci on the W chromosome p. 17
- **Table S3.** Performance statistics for the three genome assemblers p. 18
- **Table S4.** Characterization of the 50 total *P. mannii* and *P. rapae* individuals used for RAD sequencing p. 19

**Methods S1**

Details on DNA extraction and PacBio sequencing

Fresh *Pieris mannii* individuals were obtained from a single wild adult female (specimen Pi0052 in the collection of D. Berner) captured in April 2021 in Pratteln, Switzerland (site coordinates given in Table S4). This female was kept indoors at room temperature, but exposed to direct sunlight, in a 5 L glass jar with an artificial nectar source (honey-water mix, replaced daily) and host plants for oviposition (*Diplotaxis tenuifolia* and *Iberis sempervirens*, both known natural host plants of the species). Eggs produced by the female proved viable (hence the animal was already fertilized naturally upon capture) and were allowed to hatch, and the larvae were raised on fresh leaves of the same food plants. A subset of the larvae was frozen at -20°C when reaching the prepupal stage, during which no more food intake occurs while the body still remains weakly sclerotized. The other subset was allowed to develop through the pupal stage and imaginal hatch. A subset of these fresh adults was directly frozen at -20°C after hatch, but approximately a dozen of individuals were transferred for several days to outdoor aeraria (60 x 60 x 90 cm) exposed to sunlight and provided with nectar sources and host plants to attempt inbreeding. However, no full-sib crossing could be achieved; hence we decided to continue with outbred individuals from the above indoor cohort. To assess a potential effect of the preservation method on DNA integrity, both the prepupae and adults killed by freezing were partly preserved as frozen specimens, partly transferred to absolute EtOH for dehydration immediately after freezing and preserved in absolute EtOH at -20°C.

To obtain DNA of highest purity and integrity (i.e., high molecular weight), we ran test extractions using the Qiagen DNeasy Blood & Tissue (spin-column) kit, the BeckmanCoulter DNAdvance (magnetic beads) kit, and a phenol-chloroform protocol (see below) (n = 12 individuals per method). The input material further included two different tissue types: the head plus the first two (legged) segments of the prepupa, and the entire thorax of the adult (both approximately 20 – 30 mg of tissue). DNA elution was in 100 µL for all methods. The products obtained from these test extractions were then assessed for DNA purity, concentration and fragment length by combining Nanodrop One, Qubit 2.0 (broad range), TapeStation 2200 (A.02.01 SR1; genomic DNA ScreenTape), and the Agilent Femto Pulse system (software v1.0.0.32). This series of extractions revealed that all methods produced satisfactory DNA yields, but phenol-chloroform extraction clearly produced longer DNA fragments (higher integrity). Moreover, although DNA yield was slightly higher with prepupal tissue than with thorax, DNA purity tended to be higher and more consistent with the latter. No effect of the preservation method was observed, hence storing specimens frozen or in absolute EtOH can both be recommended. Based on the insights from the test extractions, we made slight optimizations to the phenol-chloroform protocol to further reduce DNA fragmentation. The final extraction protocol is presented below. To obtain DNA for sequencing, thorax tissue from three individuals per sex was subject to extraction, and based on Femto Pulse readings the best individual per sex was chosen. Gel electrophoresis and fragment length distribution images for these two individuals are displayed in fig. S8.

The DNA sample from the selected male and female were submitted to the Functional Genomics Center Zurich, Switzerland, where a Low Input 10-12 Kb HiFi library was prepared for each individual. The two libraries were then each sequenced on a full 8M SMRT Cell of a PacBio Sequel IIe instrument in HiFi/CCS mode (30 h movie). The resulting HiFi read length distributions and the read depth distributions for the v1 assemblies are presented in fig. S9.

Final version of the phenol-chloroform DNA extraction protocol, as used for the two individuals subjected to long-read sequencing

This protocol is a slight modification of the protocol provided by PacBio (https://www.pacb.com/wp-content/uploads/2015/09/SharedProtocol-Extracting-DNA-usinig-Phenol-Chloroform.pdf. Accessed 1. April 2021). All mixing steps are performed by gently inverting the tubes. All pipetting is done by using wide-bore pipette tips. For precipitation, ice-cold ethanol was used. Unless noted otherwise, reagents correspond to those described in the above PacBio protocol.

1. Cut tissue into small pieces, and place in a 1.5 ml microcentrifuge tube (this tube is called *tube 1*). Add 180 µl Buffer ATL (from Qiagen DNeasy Blood & Tissue kit). Disrupt tissue by grinding with a pestle
2. Add 20 µL Proteinase K. Mix, quick spin, and digest at 56°C during 2 h by mixing for 2 min every 30 min
3. Add 4 µL of RNase A (100 mg/mL), incubate for 15 min at room temperature
4. If necessary, bring the volume in *tube 1* to 200 µL using Elution Buffer (EB)
5. Add to *tube 1* a volume of phenol/chloroform/isoamyl alcohol (PCIA) solution equal to the current volume in *tube 1* (i.e., 200 µL)
6. Mix for 5 min
7. Spin *tube 1* at high speed for 5 min
8. Transfer ~180 µL of the supernatant aqueous solution into a new tube (*tube 2*). Avoid transferring any of the PCIA phase
9. Add 200 µL of EB to *tube 1*
10. Mix solution in *tube 1* for 5 minutes
11. Spin *tube 1* at high speed for 5 min
12. Transfer as much of the supernatant aqueous solution as possible from *tube 1* into *tube 2*, avoiding the transfer of any of the PCIA phase. Discard *tube 1*
13. Add to *tube 2* a volume of chloroform/isoamyl alcohol (CIA) solution equal to the current volume in *tube 2*
14. Mix solution in *tube 2* for 5 min
15. Spin *tube 2* at high speed for 5 min
16. Transfer as much of the supernatant aqueous solution as possible from *tube 2* into a new tube (*tube 3*), avoiding the transfer of any of the CIA phase. Discard *tube 2*
17. Add to *tube 3* $NH_4OAc$ to a final concentration of 0.75 M
18. Mix solution for 5 min
19. To precipitate, add to *tube 3* a volume of 100% ethanol 2.5x the current volume in *tube 3*, and mix for 5 min
20. Incubate at -20°C for 30 min
21. Spin for 20 min at top speed in a 4°C centrifuge
22. Decant and discard the liquid carefully without disrupting the pellet on the tube bottom
23. Wash the pellet by adding 300 µL of 80% EtOH and mixing for 5 min

24. Spin for 15 min at top speed in a 4°C centrifuge
25. Decant and discard the liquid carefully without disrupting the pellet on the tube bottom
26. For a second 80% EtOH wash, repeat steps 23-25
27. Quick spin on table top centrifuge to pull residual EtOH to the bottom
28. Remove residual EtOH with a P20 pipette, without disrupting the pellet
29. Air dry for 1-2 min
30. Re-suspend in desired volume of EB (85 µL was used)

**Fig. S1**

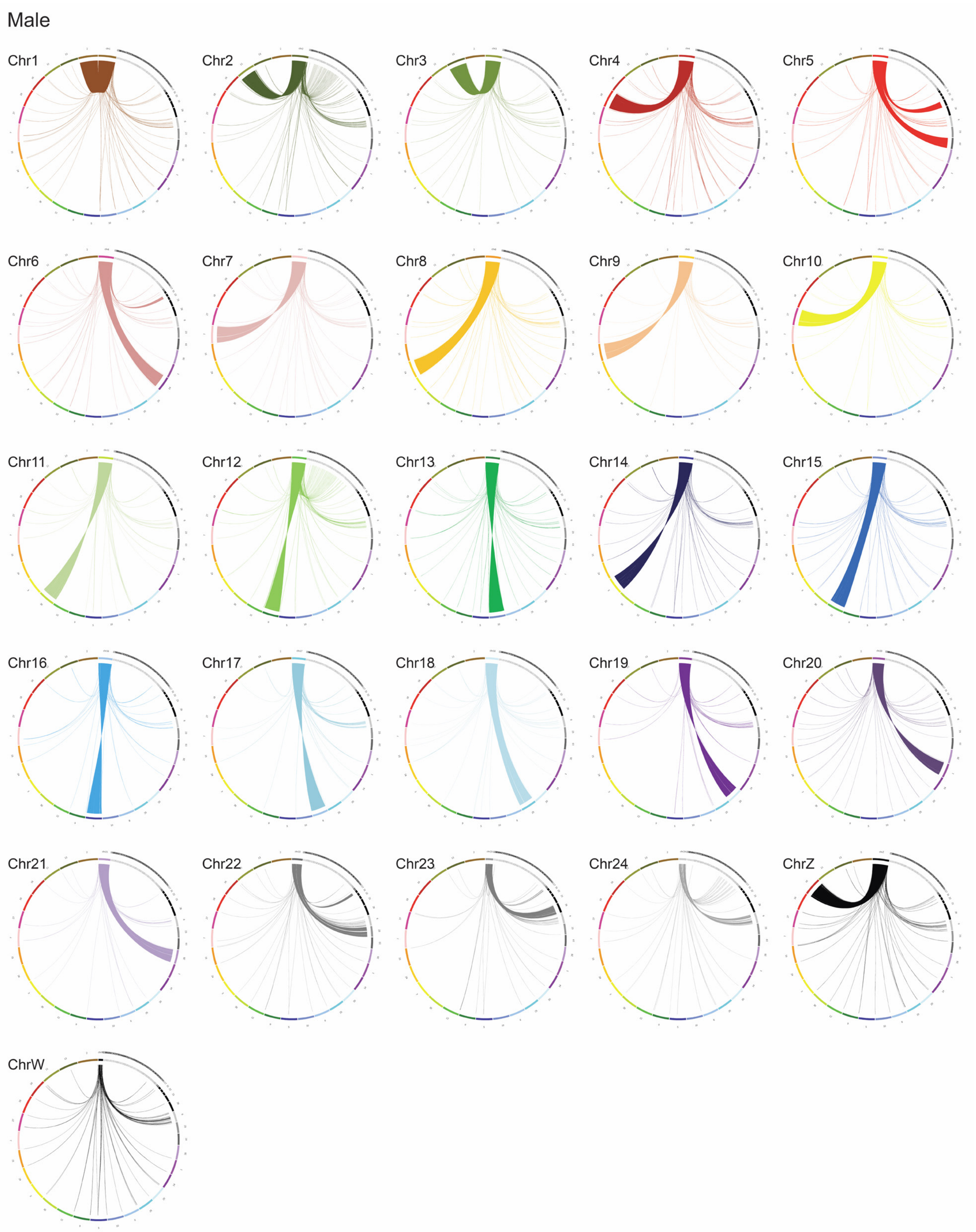
Male
Chr1
Chr2
Chr3
Chr4
Chr5
Chr6
Chr7
Chr8
Chr9
Chr10
Chr11
Chr12
Chr13
Chr14
Chr15
Chr16
Chr17
Chr18
Chr19
Chr20
Chr21
Chr22
Chr23
Chr24
ChrZ
ChrW

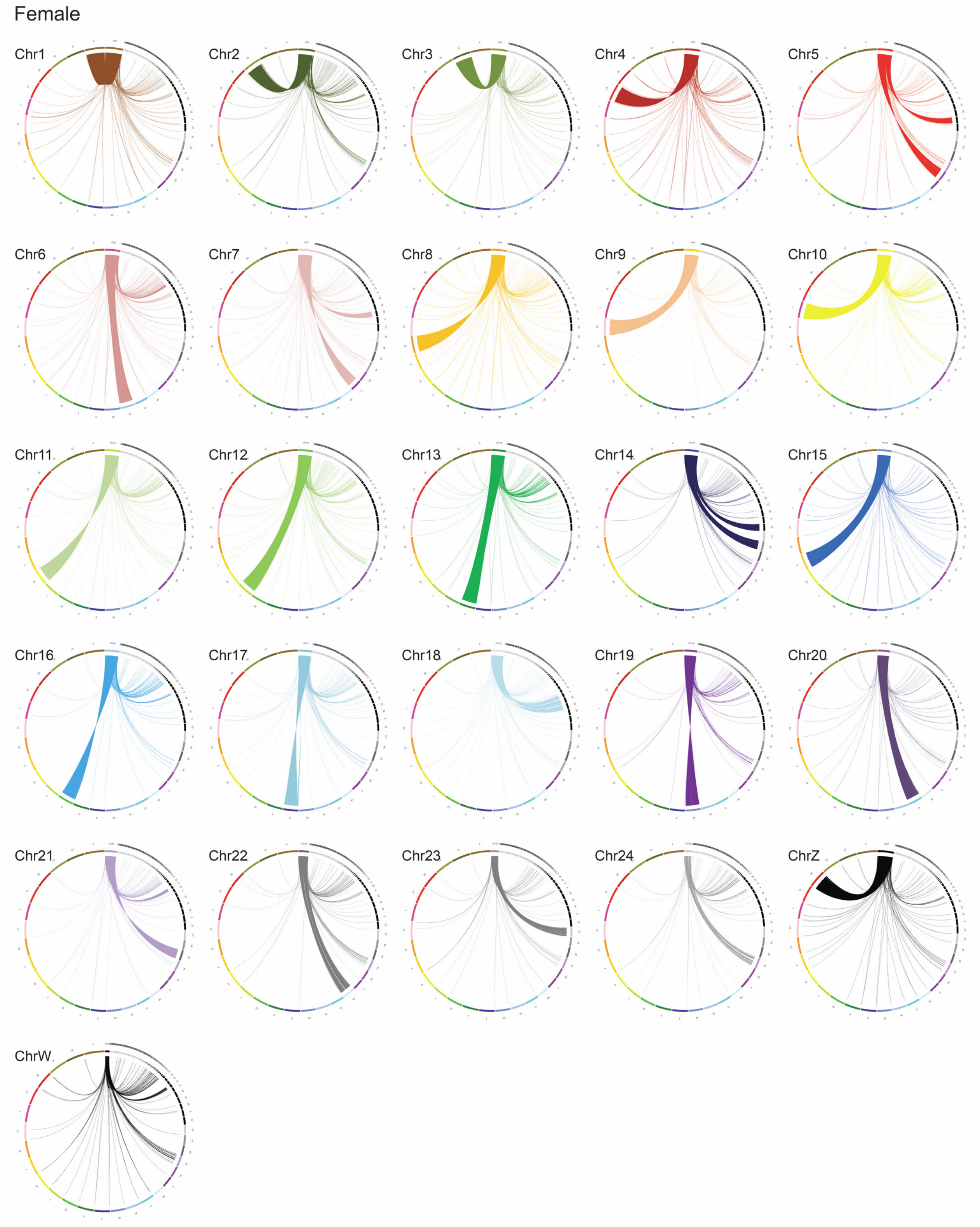


**Fig. S1.** Alignment of individual *P. rapae* chromosomes to the *P. mannii* male and female v1 assemblies using Minimap2. For the female v2 assembly, contig 37, representing the W chromosome, was replaced by a slightly longer contig obtained by an alternative assembly considering only the HiFi reads longer than 5 kb. The relationship between the *P. rapae* chromosomes and *P. mannii* contigs is also summarized in table S1.

### Fig. S2

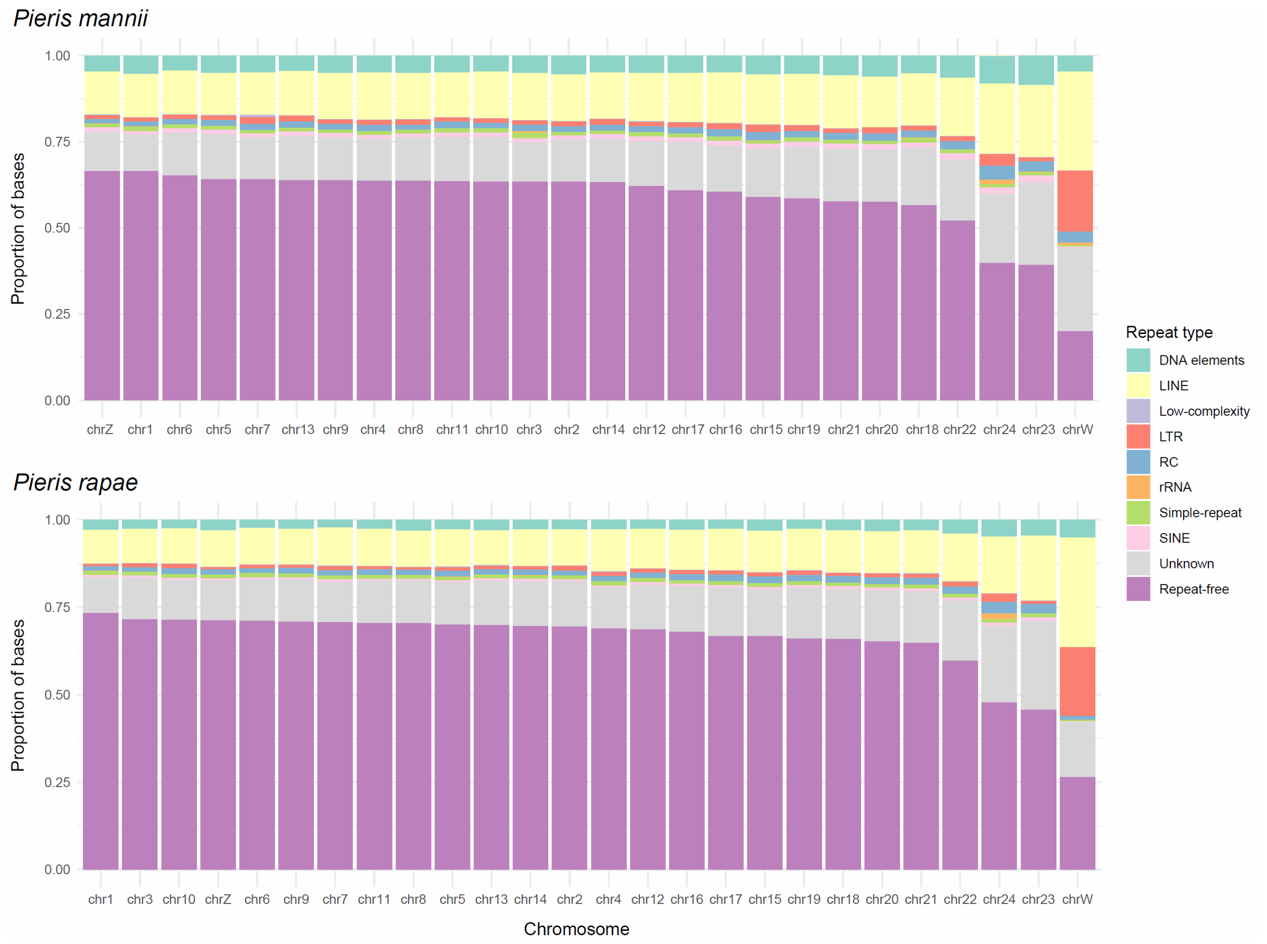


**Fig. S2.** Proportion of each chromosome of *P. mannii* and *P. rapae* assigned to different repeat classes by RepeatMasker. The category on the bottom (purple) indicates the proportion of non-repeated DNA.

**Fig. S3**

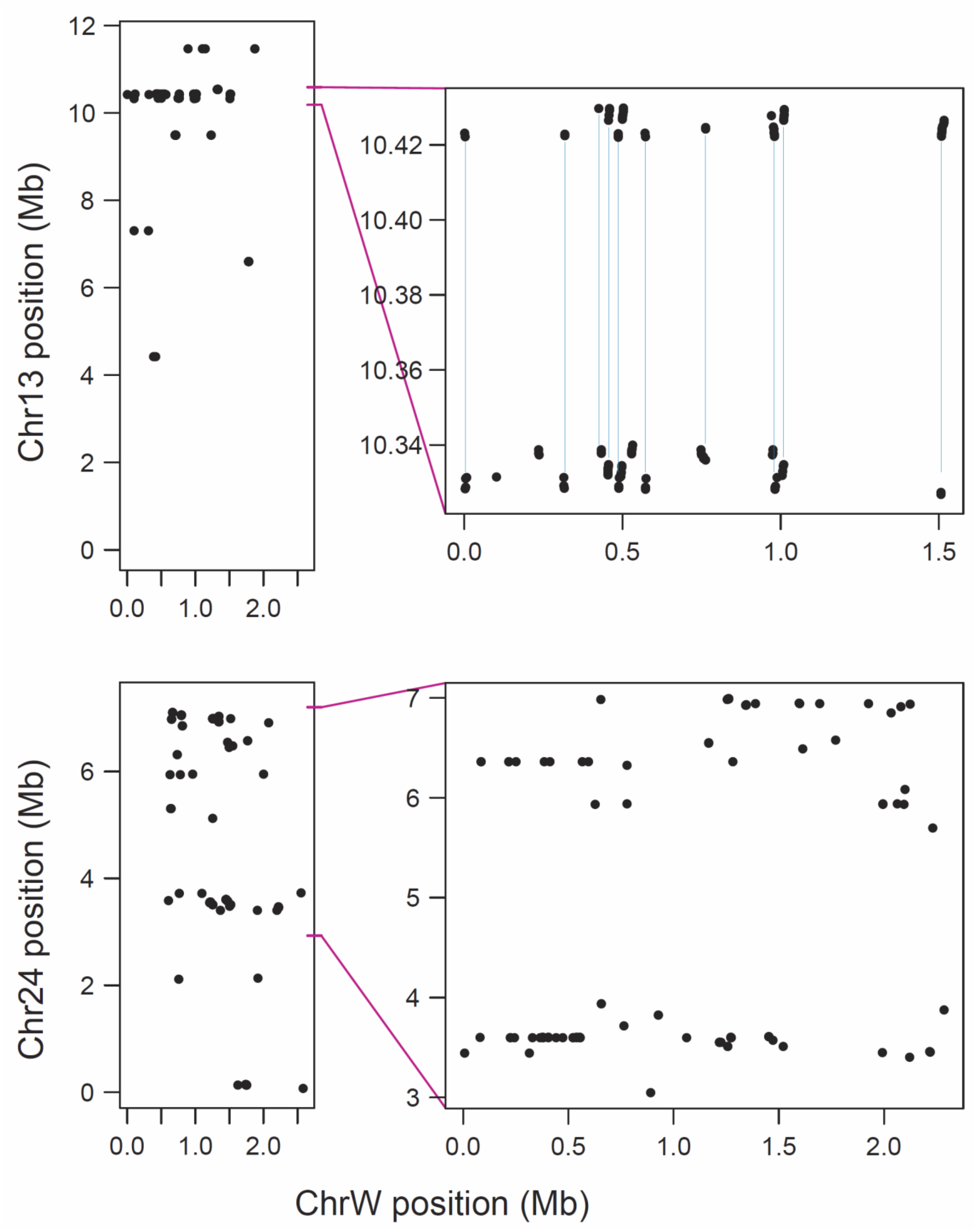


**Fig. S3.** Physical position of *P. mannii* W chromosome sequence tags aligning to the chromosomes 13 (top) and 24 (bottom), revealing extensive repeat expansion of autosomal sequences on the W. As in fig. 7, the left graphics are based on sequence tags sampled from the repeat-masked W chromosome, while the right graphs represent tags from the W not repeat-masked. The upper close-up indicates extensive repeat expansion on the W of DNA from two regions on chromosome 13 (51% of the W sequence tags aligning to this autosome map to these two regions). Segments from these regions appear to be repeated in tandem, as highlighted by the vertical lines, thus suggesting joint copy multiplication.

**Fig. S4**

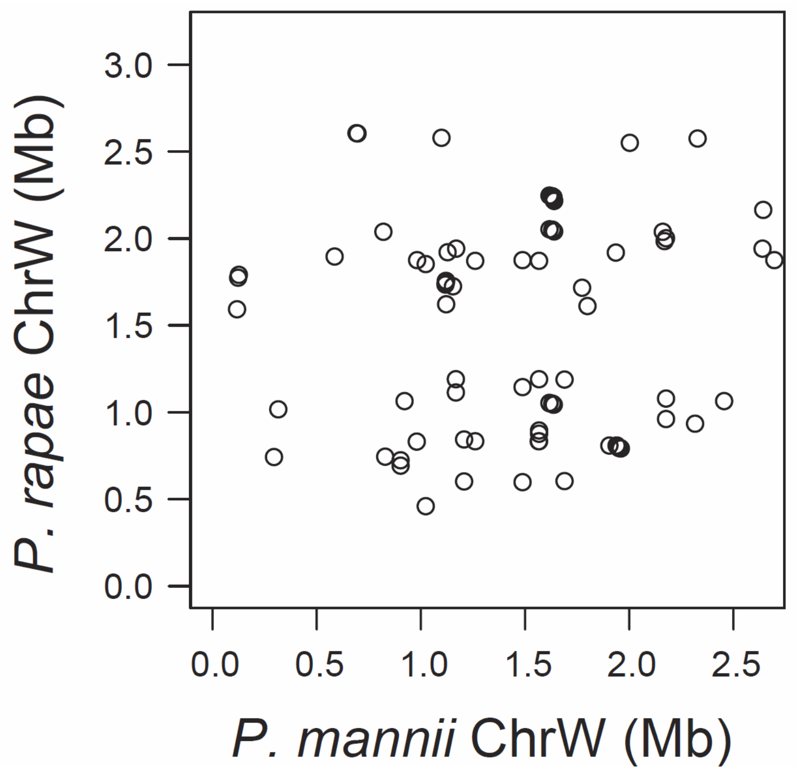


**Fig. S4.** Positions of sequence tags from the *P. mannii* W chromosome plotted against their corresponding alignment positions on the *P. rapae* W chromosome, as in fig. 2C. The difference is that here the sequence tags are derived from the repeat-masked W (n = 203 tags with unique alignment). Both analyses consistently indicate massive divergence in W chromosome structure between the sister species.

**Fig. S5**

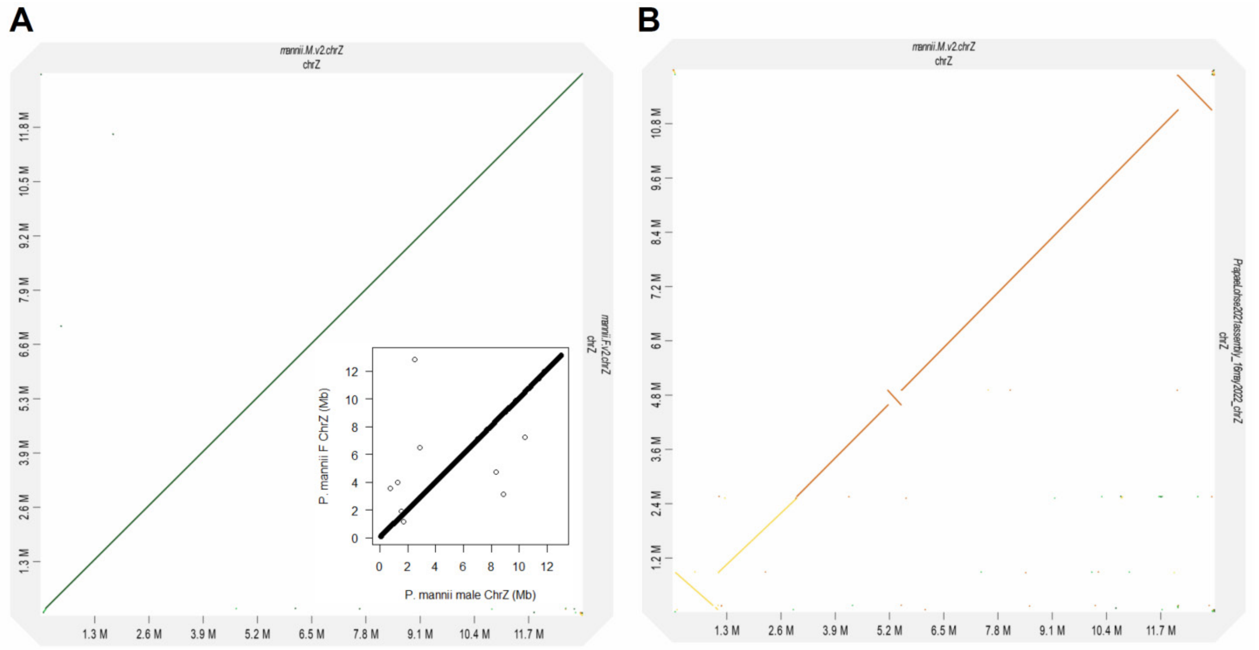


**Fig. S5** Check for correct assembly of the *P. mannii* Z chromosomes. (A) Alignment of the female to the male *P. mannii* Z chromosome, confirming perfect collinearity along the entire assemblies. (B) Alignment of the *P. rapae* to the male *P. mannii* Z chromosome. Apart from three minor inconsistencies in chromosome segment orientation, the Z chromosome builds of the two sister species are also collinear. Because the two *P. mannii* Z assemblies were performed absolutely independently and do not display any conflict in orientation, the between-species inconsistencies must either reflect inversion polymorphisms, or, more plausibly, assembly errors within *P. rapae*. The main plots were generated using D-GENIES, based on whole-chromosome Minimap2 alignment. The insert in (A) is based on 24,177 sequence tags extracted from the male Z chromosome and aligned to the female genome (both not repeat-masked).

**Fig. S6**

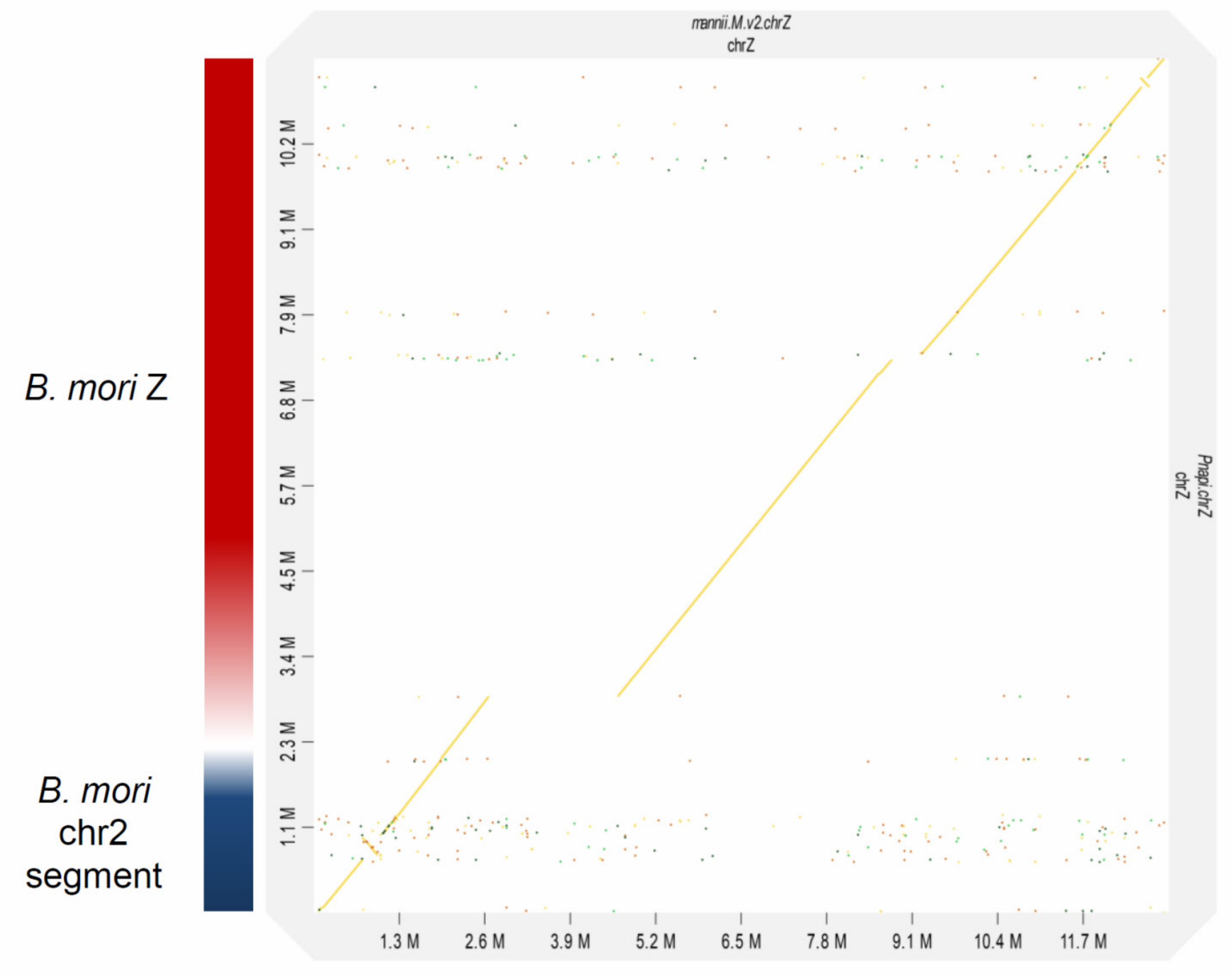


**Fig. S6** Alignment of the *Pieris napi* to the *P. mannii* Z chromosome. Apart from putative assembly gaps in the *P. napi* Z assembly, the Z chromosome builds are highly collinear between the species. According to Hill et al. (2019), the proximal chromosome segment of the *P. napi* Z (= chr1) originates from a fusion between the ancestral lepidopteran Z chromosome (chromosome 1 in the *Bombyx mori* reference genome; shown as red bar on the left of the graphic), and a segment from the autosome corresponding to chromosome 2 in the *B. mori* genome (shown as blue bar). This latter segment is also located on the proximal ('left') side in the *P. mannii* Z assembly. The plot was generated using D-GENIES based on Minimap2 alignment.

**Fig. S7**

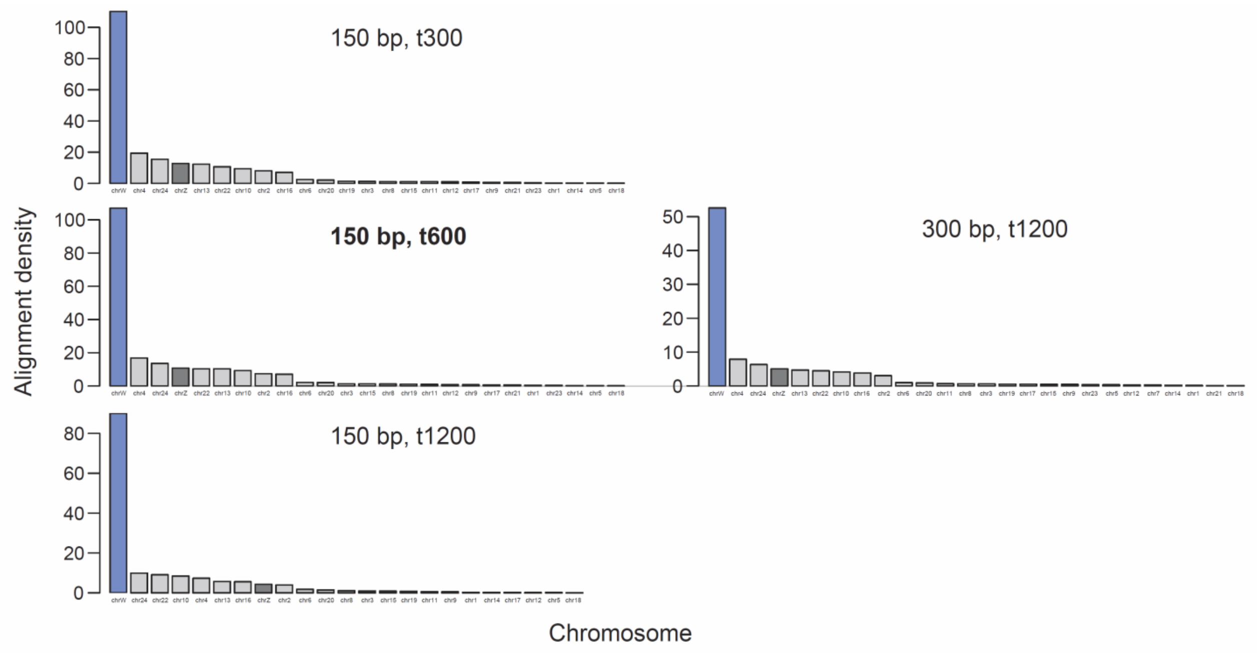


**Fig. S7** Robustness check evaluating the effect of different total alignment mismatch thresholds (t-flag in Novoalign; t300, 600, 1200) and different DNA segment lengths (150 and 300 bp) on the alignment density of *P. mannii* W chromosome tags on *P. rapae* chromosomes. In bold are the default settings underlying the analyses presented in the paper. Although both mismatch threshold and segment length influence the absolute alignment success and hence the Y-axis scale, relative differences in alignment density among chromosomes are highly consistent across the settings evaluated. Note that with 300 bp, a mismatch of t1200 results in the same per-base mismatch stringency as t600 with 150 bp. Also, although alignment success per read was slightly higher with 300 bp than with 150 bp, the alignment density is lower for 300 bp because fewer of the longer tags could be extracted from the repeat-masked W chromosome.

**Fig. S8**

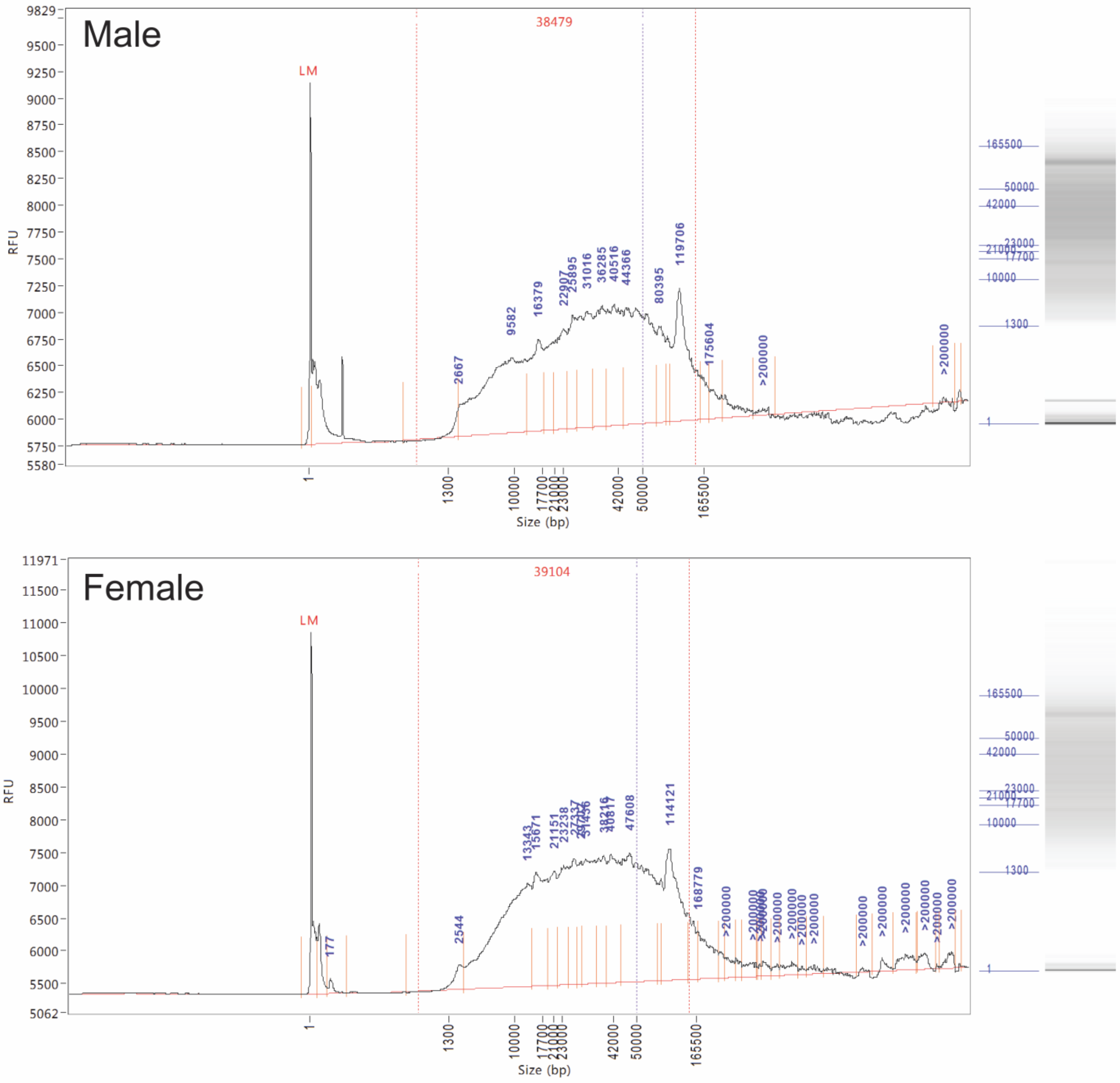


**Fig. S8.** Fragment length density distribution (left) and gel electrophoresis image (right) for the DNA extracted with the phenol-chloroform protocol (Methods S1) from the male and female *P. mannii* individual, as quantified by Femto Pulse. The red number on the top of each graph indicates mean fragment length, which is close to 40 kb for both individuals.

**Fig. S9**

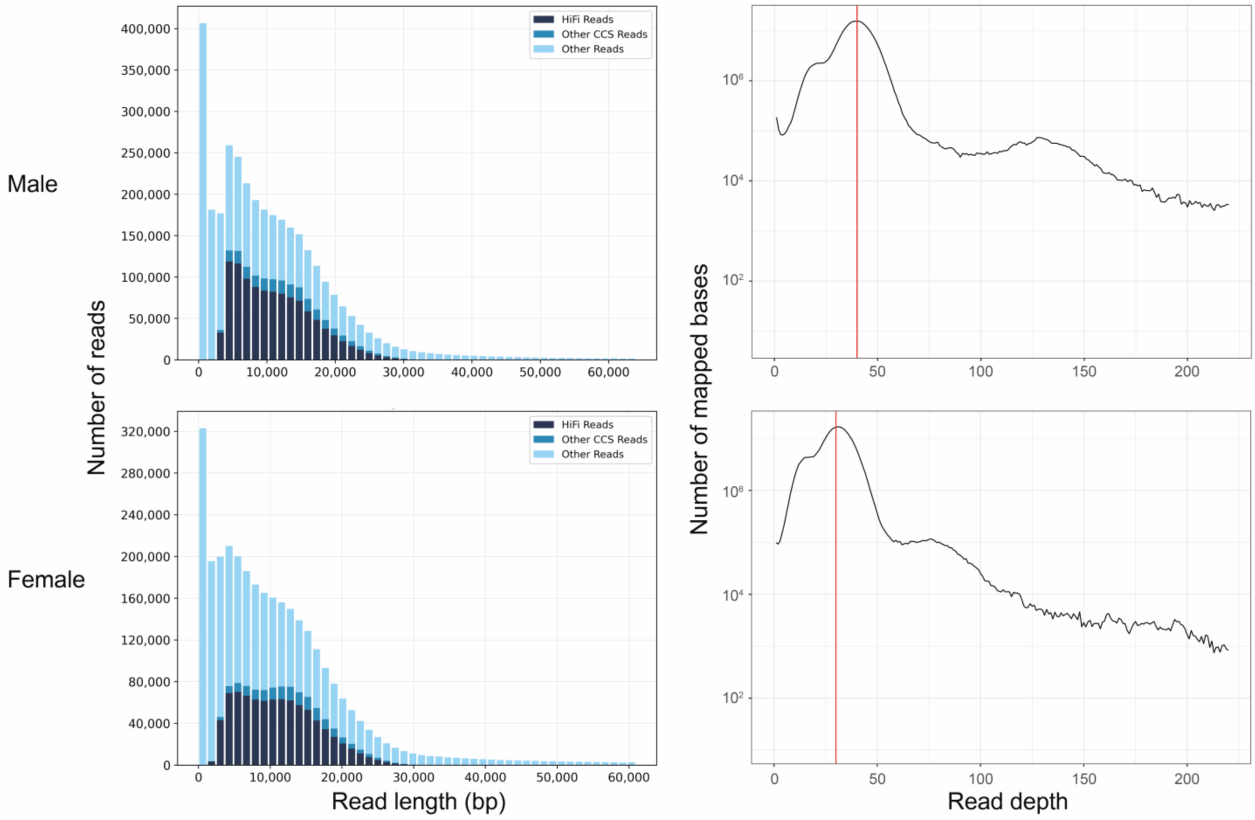


**Fig. S9.** Length distribution of the reads generated by PacBio long-read sequencing of the male and female *P. mannii* individuals (left column). Only the HiFi reads were used for genome assembly. The right column shows the density distribution of HiFi read depth per base position, as quantified by mapping the HiFi reads to the v1 genome assemblies.

**Table S1**. Characterization of the male and female *P. mannii* contigs homologous to *P. rapae* chromosomes, as obtained from the Hifiasm v1 assemblies. The chromosome (v2) numbers follow the designation used in the *P. rapae* reference genome. Contigs marked by an asterisk were assembled in reverse orientation relative to the corresponding chromosome in the sister species, and hence were reverse-complemented for the v2 build. The length column gives the final v2 chromosome length, hence in a few cases combines multiple contigs. For the W chromosome, the length indicated refers to a slightly longer assembly obtained by considering only the HiFi reads longer than 5 kb (length of the original v1 contig 37: 2'416'388). The telomere column specifies separately for each chromosome end whether the insect-specific (TTAGG)n telomere repeats are present.

| | Male assembly | | | Female assembly | | |
|---|---|---|---|---|---|---|
| Chromosome (v2) | Contigs (v1) | Length (bp) | Telomere | Contigs (v1) | Length (bp) | Telomere |
| 1 | 2 | 14'165'569 | y/y | 6 | 14'525'974 | y/y |
| 2 | 11 | 13'263'370 | y/y | 16 | 13'294'971 | y/y |
| 3 | 13 | 13'443'999 | y/y | 12* | 13'552'510 | y/y |
| 4 | 18* | 12'768'226 | y/y | 13 | 12'690'870 | y/y |
| 5 | 19*, 21* | 12'604'376 | y/y | 29, 5 | 12'617'736 | y/y |
| 6 | 14*, 31 | 12'479'271 | y/y | 18*, 17 | 12'398'157 | y/y |
| 7 | 7 | 12'635'611 | y/y | 2, 26 | 12'383'211 | y/y |
| 8 | 24* | 12'354'463 | y/y | 10 | 12'424'943 | y/y |
| 9 | 15 | 12'512'088 | y/y | 21* | 12'471'614 | y/y |
| 10 | 27* | 12'656'134 | y/y | 7* | 12'501'273 | y/y |
| 11 | 30 | 12'171'410 | y/y | 19 | 12'173'198 | y/y |
| 12 | 9 | 11'767'063 | y/y | 1* | 11'803'600 | n/y |
| 13 | 10 | 11'628'137 | y/y | 9* | 11'707'024 | y/y |
| 14 | 3 | 12'214'670 | y/y | 34*, 22* | 12'194'447 | y/y |
| 15 | 12* | 12'121'942 | y/y | 3* | 12'183'372 | y/y |
| 16 | 5 | 11'685'795 | y/y | 25 | 11'792'990 | y/y |
| 17 | 8 | 11'242'583 | y/y | 8 | 11'205'745 | y/n |
| 18 | 25* | 11'103'484 | y/y | 32, 24*, 4 | 11'223'805 | y/y |
| 19 | 20 | 10'818'668 | y/y | 30 | 10'764'362 | y/y |
| 20 | 1 | 10'319'200 | y/y | 11* | 10'295'211 | n/y |
| 21 | 28* | 10'025'374 | y/y | 20, 27* | 9'848'088 | y/y |
| 22 | 23*, 17 | 9'448'146 | y/y | 40* | 9'182'131 | y/y |
| 23 | 26* | 7'027'187 | y/y | 15* | 6'587'828 | n/y |
| 24 | 6 | 7'370'422 | y/y | 14* | 7'659'231 | y/y |
| Z | 16* | 13'053'618 | y/y | 28* | 13'155'917 | n/y |
| W | | | | 37 | 2'639'529 | n/y |
| Mitochondrial | | 15'163 | | | 15'158 | |

**Table S2**. Female-limited RAD loci on the W chromosome. These 13 RAD loci were present in all 19 females but in none of the 19 males. The first position ('residual') specifies the 5-prime W chromosome start position of the Pst1 restriction enzyme residual (TGCAG) relative to the strand to which the short reads effectively aligned (indicated in the third column). The second position ('+ strand') reports the standard 5-prime start position on the + strand for all loci. The sequences represent the exact haplotype observed most frequently among the 19 females ('major haplotype'). The number of females exhibiting this specific haplotype is indicated in the last column.

| Position (residual) | Position (+ strand) | Strand | Sequence (major haplotype) | Frequency |
|---|---|---|---|---|
| 700'639 | 700'549 | - | TGCAGAACTGGCCAAATCTCCTCTCTTACGTCTGGGCATTGTATTCTATAAAATATAAATTAAATAAACATTTTATAGATAATTATTTTGT | 18 |
| 736'825 | 736'735 | - | TGCAGAGCACATTTTTTTTTTCTTTTATAACATTTTTCGAAATATTATTTTGCTTTAATCTGCCCTTGTAATATTTTATTAATATTTTTAC | 11 |
| 792'223 | 792'223 | + | TGCAGATTTTTTGTGTAGACAGTTGTAAGTTTATTTGATTTGTTTTCCTTAACTCAAAACAGGCGGGTGTAAGGCCATCAATAAATTCTTT | 14 |
| 894'943 | 894'943 | + | TGCAGATAATACCTAATCTAATATTAACAATATTATACATGTTTTAAGATTTTTATGACTTCTCACCAATATAAGGATATATAAGTATCTA | 17 |
| 1'235'425 | 1'235'335 | - | TGCAGCGCGAGAATTAGGTTCATTTATCAAACAAAATAAAGATTTTCTGTTTGACTTTGCAATTCCAATATATTTCATTTCATTCCGGCAT | 18 |
| 1'255'344 | 1'255'344 | + | TGCAGCAAGGGTGGAGGGGCTGCGTTTTCAACAACAACTTTAAACAGAACAATCTTATTAAACTGGTTCCTTACAGTTTCCTTTCCTAAAT | 17 |
| 1'331'719 | 1'331'629 | - | TGCAGAAATTGCTCACGTTGTCCTCCATTTTCGCGGATTGTTTGTGTTGAGTTTTGTTCTTTTCAATCTTCTCACTAGGCCCGTGGTGTCA | 18 |
| 1'714'917 | 1'714'917 | + | TGCAGAACCGGCCAAATCTCCTCCTCTCTTACGTCTGGGCATTGTATTCTATAAAATATAAATTAAATAAACATTTTATAGATAATTATTT | 18 |
| 1'760'139 | 1'760'139 | + | TGCAGCACACATAATTAATACACTGGCTAATAGTTATAATTTATTTTAAGAATATTCATAATTAATATTTTCGTGGCAAACAAATTGAAAA | 17 |
| 1'810'058 | 1'810'058 | + | TGCAGATGCAGTGAAACGTTTTGACATTCCCACTGAGCATCCAACCGAGCTTAGTTTGTTGCGCAGTCGGCAGATTTGCATTTGTTCTACA | 19 |
| 1'845'716 | 1'845'716 | + | TGCAGAGCACATTTTTTTTTCTTTTATAACATTTTTCGAAATATTTGTTTGCTTTAATCTGCCCTTGTAATATTTTATTAATATTTTTACA | 10 |
| 1'897'796 | 1'897'706 | - | TGCAGTCCATTTTGTTGCAGCATTAAAGACGGCGGTTGTGACAGGCTTAACTGCACCACATAGCCGAGGGGCCAACTGTGAAGCAGATGAT | 17 |
| 2'561'313 | 2'561'313 | + | TGCAGCCTGGCGCTTTCTTCGTGTAGTATGTCCACAGAACATTCAGATCGGTTACGGTTGTTTGTCGCTAAGACCATAAGTACTTGTACAC | 17 |

| **Table S3**. Comparison of the performance of the three long-read assemblers considered for the raw assembly of the male and female *P. mannii* genomes. The performance statistics were obtained from QUAST and BUSCO (last row). The BUSCO values refer to the complete gene category obtained with the Lepidoptera database (n = 5286 genes). | | | | | | |
|---|---|---|---|---|---|---|
| | Male v1 assembly | | | Female v1 assembly | | |
| Assembly software | Hifiasm | HiCanu | IPA | Hifiasm | HiCanu | IPA |
| Number of contigs | 111 | 160 | 187 | 103 | 134 | 182 |
| Longest contig (Mb) | 14 | 13.5 | 10.5 | 14.5 | 12.5 | 10.5 |
| Total assembly length (Mb) | 296 | 297 | 287 | 303 | 297 | 290 |
| N50 (Mb) | 12.1 | 9 | 4.5 | 11.8 | 5.8 | 3.8 |
| L50 (Mb) | 12 | 14 | 22 | 12 | 17 | 24 |
| BUSCO % complete | 99.2 | 99.1 | 98.5 | 99.0 | 99.4 | 97.9 |

**Table S4**. Individuals of *P. mannii* and *P. rapae* used for RAD sequencing. The last column specifies the accession numbers to the individual short read data in the NCBI Short Read Archive.

| ID_photo | ID_RADseq | Species | Sex | Sampling location | Latitude | Longitude | Year sampled | NCBI accession number |
|---|---|---|---|---|---|---|---|---|
| Pi0001 | CH_Prt_BeG_01 | *mannii* | f | Pratteln | 47.52069 | 7.68656 | 2020 | SRR21859196 |
| Pi0002 | CH_Prt_BeG_02 | *mannii* | m | Pratteln | 47.52069 | 7.68656 | 2020 | SRR21859195 |
| Pi0005 | CH_Prt_BeG_03 | *mannii* | m | Pratteln | 47.52069 | 7.68656 | 2020 | SRR21859184 |
| Pi0006 | CH_Prt_BeG_04 | *mannii* | m | Pratteln | 47.52069 | 7.68656 | 2020 | SRR21859173 |
| Pi0007 | CH_Prt_BeG_05 | *mannii* | f | Pratteln | 47.52069 | 7.68656 | 2020 | SRR21859162 |
| Pi0008 | CH_Prt_BeG_06 | *mannii* | m | Pratteln | 47.52069 | 7.68656 | 2020 | SRR21859151 |
| Pi0009 | CH_Prt_BeG_07 | *mannii* | m | Pratteln | 47.52069 | 7.68656 | 2020 | SRR21859150 |
| Pi0012 | CH_Prt_BeG_10 | *mannii* | m | Pratteln | 47.52069 | 7.68656 | 2020 | SRR21859149 |
| Pi0013 | CH_Prt_BeG_11 | *mannii* | f | Pratteln | 47.52069 | 7.68656 | 2020 | SRR21859148 |
| Pi0014 | CH_Prt_BeG_12 | *mannii* | m | Pratteln | 47.52069 | 7.68656 | 2020 | SRR21859147 |
| Pi0015 | CH_Prt_BeG_13 | *mannii* | f | Pratteln | 47.52069 | 7.68656 | 2020 | SRR21859194 |
| Pi0017 | CH_Prt_BeG_14 | *mannii* | m | Pratteln | 47.52069 | 7.68656 | 2020 | SRR21859193 |
| Pi0018 | CH_Prt_BeG_15 | *mannii* | m | Pratteln | 47.52069 | 7.68656 | 2020 | SRR21859192 |
| Pi0022 | CH_Prt_BeG_18 | *mannii* | f | Pratteln | 47.52069 | 7.68656 | 2020 | SRR21859191 |
| Pi0023 | CH_Prt_BeG_19 | *mannii* | f | Pratteln | 47.52069 | 7.68656 | 2020 | SRR21859190 |
| Pi0024 | CH_Prt_BeG_20 | *mannii* | f | Pratteln | 47.52069 | 7.68656 | 2020 | SRR21859189 |
| Pi0025 | CH_Prt_BeG_21 | *mannii* | m | Pratteln | 47.52069 | 7.68656 | 2020 | SRR21859188 |
| Pi0029 | CH_Prt_BeG_25 | *mannii* | f | Pratteln | 47.52069 | 7.68656 | 2020 | SRR21859187 |
| Pi0030 | CH_Prt_BeG_26 | *mannii* | f | Pratteln | 47.52069 | 7.68656 | 2020 | SRR21859186 |
| Pi0034 | CH_Prt_BeG_28 | *mannii* | m | Pratteln | 47.52069 | 7.68656 | 2020 | SRR21859185 |
| Pi0037 | CH_Prt_BeG_31 | *mannii* | f | Pratteln | 47.52069 | 7.68656 | 2020 | SRR21859183 |
| Pi0042 | CH_Prt_BeG_35 | *mannii* | f | Pratteln | 47.52069 | 7.68656 | 2020 | SRR21859182 |
| Pi0044 | CH_Prt_BeG_37 | *mannii* | f | Pratteln | 47.52069 | 7.68656 | 2020 | SRR21859181 |
| Pi0046 | CH_Prt_BeG_39 | *mannii* | m | Pratteln | 47.52069 | 7.68656 | 2020 | SRR21859180 |
| Pi0087 | CH_Bsl_Zie_01 | *mannii* | m | Ziefen | 47.43230 | 7.70540 | 2021 | SRR21859179 |
| Pi0088 | CH_Bsl_Zie_02 | *mannii* | m | Ziefen | 47.43230 | 7.70540 | 2021 | SRR21859178 |
| Pi0089 | CH_Bsl_Zie_03 | *mannii* | f | Ziefen | 47.43230 | 7.70540 | 2021 | SRR21859177 |
| Pi0092 | CH_Bsl_Gri_01 | *mannii* | f | Grindel | 47.38047 | 7.50421 | 2021 | SRR21859176 |
| Pi0093 | CH_Bsl_Gri_02 | *mannii* | m | Grindel | 47.38047 | 7.50421 | 2021 | SRR21859175 |
| Pi0094 | CH_Bsl_Zie_04 | *mannii* | m | Ziefen | 47.43230 | 7.70540 | 2021 | SRR21859174 |
| Pi0096 | CH_Bsl_Zie_05 | *mannii* | f | Ziefen | 47.43230 | 7.70540 | 2021 | SRR21859172 |
| Pi0110 | CH_Bsl_Bsl_04 | *mannii* | f | Basel | 47.56717 | 7.56740 | 2021 | SRR21859171 |
| Pi0137 | CH_Bsl_Pfe_01 | *mannii* | f | Pfeffingen | 47.45749 | 7.59467 | 2021 | SRR21859170 |
| Pi0138 | CH_Bsl_Pfe_02 | *mannii* | m | Pfeffingen | 47.45749 | 7.59467 | 2021 | SRR21859169 |
| Pi0139 | CH_Bsl_Pfe_03 | *mannii* | f | Pfeffingen | 47.45749 | 7.59467 | 2021 | SRR21859168 |
| Pi0140 | CH_Bsl_Pfe_04 | *mannii* | m | Pfeffingen | 47.45749 | 7.59467 | 2021 | SRR21859167 |

| | | | | | | | | |
|---|---|---|---|---|---|---|---|---|
| Pi0141 | CH_Bsl_Pfe_05 | *mannii* | f | Pfeffingen | 47.45749 | 7.59467 | 2021 | SRR21859166 |
| Pi0142 | CH_Bsl_Pfe_06 | *mannii* | m | Pfeffingen | 47.45749 | 7.59467 | 2021 | SRR21859165 |
| Pi0019 | CH_Prt_BeG_16 | *rapae* | f | Pratteln | 47.52069 | 7.68656 | 2020 | SRR21859164 |
| Pi0021 | CH_Prt_BeG_17 | *rapae* | f | Pratteln | 47.52069 | 7.68656 | 2020 | SRR21859163 |
| Pi0027 | CH_Prt_BeG_23 | *rapae* | f | Pratteln | 47.52069 | 7.68656 | 2020 | SRR21859161 |
| Pi0028 | CH_Prt_BeG_24 | *rapae* | f | Pratteln | 47.52069 | 7.68656 | 2020 | SRR21859160 |
| Pi0032 | CH_Prt_BeG_27 | *rapae* | m | Pratteln | 47.52069 | 7.68656 | 2020 | SRR21859159 |
| Pi0036 | CH_Prt_BeG_30 | *rapae* | m | Pratteln | 47.52069 | 7.68656 | 2020 | SRR21859158 |
| Pi0041 | CH_Prt_BeG_34 | *rapae* | m | Pratteln | 47.52069 | 7.68656 | 2020 | SRR21859157 |
| Pi0047 | CH_Prt_BeG_40 | *rapae* | m | Pratteln | 47.52069 | 7.68656 | 2020 | SRR21859156 |
| Pi0059 | CH_Zue_All_01 | *rapae* | f | Zürich | 47.40143 | 8.53725 | 2021 | SRR21859155 |
| Pi0061 | CH_Zue_All_02 | *rapae* | m | Zürich | 47.40143 | 8.53725 | 2021 | SRR21859154 |
| Pi0062 | CH_Zue_All_03 | *rapae* | m | Zürich | 47.40143 | 8.53725 | 2021 | SRR21859153 |
| Pi0063 | CH_Zue_All_04 | *rapae* | f | Zürich | 47.40143 | 8.53725 | 2021 | SRR21859152 |
